# The effect of two distinct viral-enriched inocula on the immune response and chemical cues in honey bee pupae

**DOI:** 10.1101/2025.10.20.683288

**Authors:** Tal Erez, Angelina Fathia Osabutey, Elad Bonda, Assaf Otmy, Sofia Levin-Nikulin, Poppy J Hesketh-Best, Clarissa Pellegrini Ferreira, Marla Spivak, Declan C. Schroeder, Nor Chejanovsky, Victoria Soroker

## Abstract

The combination of *Varroa destructor* (Varroa) and the viruses it vectors is a major driver of honey bee (*Apis mellifera*) colony losses. Hygienic behavior and individual immunity enable bees to cope with some pests and parasites. Hygienic response of adult bees is driven by olfactory cues emanating from diseased brood. In this study, we tested the effect of viral inoculation on the chemical profile and immune response of pupae, independently of Varroa.

We injected pupae with various doses of an enriched inocula of either a deformed wing virus (DWV) B-A recombinant or Israeli acute paralysis virus (IAPV) and followed their development and survival for five days. At the end of the five-day period, volatile profiles, viral loads, and expression of seven immune genes were assessed. Both IAPV and the DWV loads increased to equivalent high levels irrespective of the initial dose applied. Notably, greater rates of mortality (60% loss) were observed with the highest IAPV dose when compared to the lowest IAPV dose (15% loss). The DWV inoculum, caused limited mortality but nonetheless inhibited pupae development. The IAPV injected pupae showed evidence of a dose dependent DWV amplification from either a DWV variant in the original IAPV inoculum or from an endogenous source of DWV in the pupae themselves. IAPV-injected pupae had lower expression of immune genes than DWV-injected pupae, suggesting IAPV inhibits the pupae immune response. Overall, among the seven tested immune genes six were upregulated with only *vago* downregulated, suggesting inhibition of the RNAi pathway following infection. Chemical cues of mock and untreated pupae were similar, but notably different from the virus-injected pupae for both inocula. Our findings show that Varroa-independent virus inoculated pupae produce unique virus-specific chemical cues; the ultimate consequence of such a change might lead to virus specific bee behavioral responses.

**Authors summary:** This study investigated how two major honey bee viruses, Deformed Wing Virus (DWV) and Israeli Acute Paralysis Virus (IAPV), affect pupae independent of the Varroa mite, which typically spreads them. This study was conducted in the laboratory on incubated pupae isolated from the colony and injected with two viral preparations (inocula). We tested the impact after five-days incubation. The results show that both viruses reach high levels in pupae, though IAPV inoculum proved far more deadly (up to 60% mortality) and appeared to suppress the pupae’s immune response, possibly even amplifying co-occurring DWV. While less lethal, the DWV inoculum still inhibited normal development. Interestigly, the virus-infected pupae produced distinct, virus-specific chemical cues—unique volatile profiles different from healthy controls. Since adult honey bees use olfactory cues to detect and remove diseased brood (hygienic behavior), these findings suggest that the viruses themselves generate the “scent of sickness,” which could trigger colony-wide behavioral responses and offer a vital target for enhancing the bees’ natural defenses against these major drivers of colony loss.

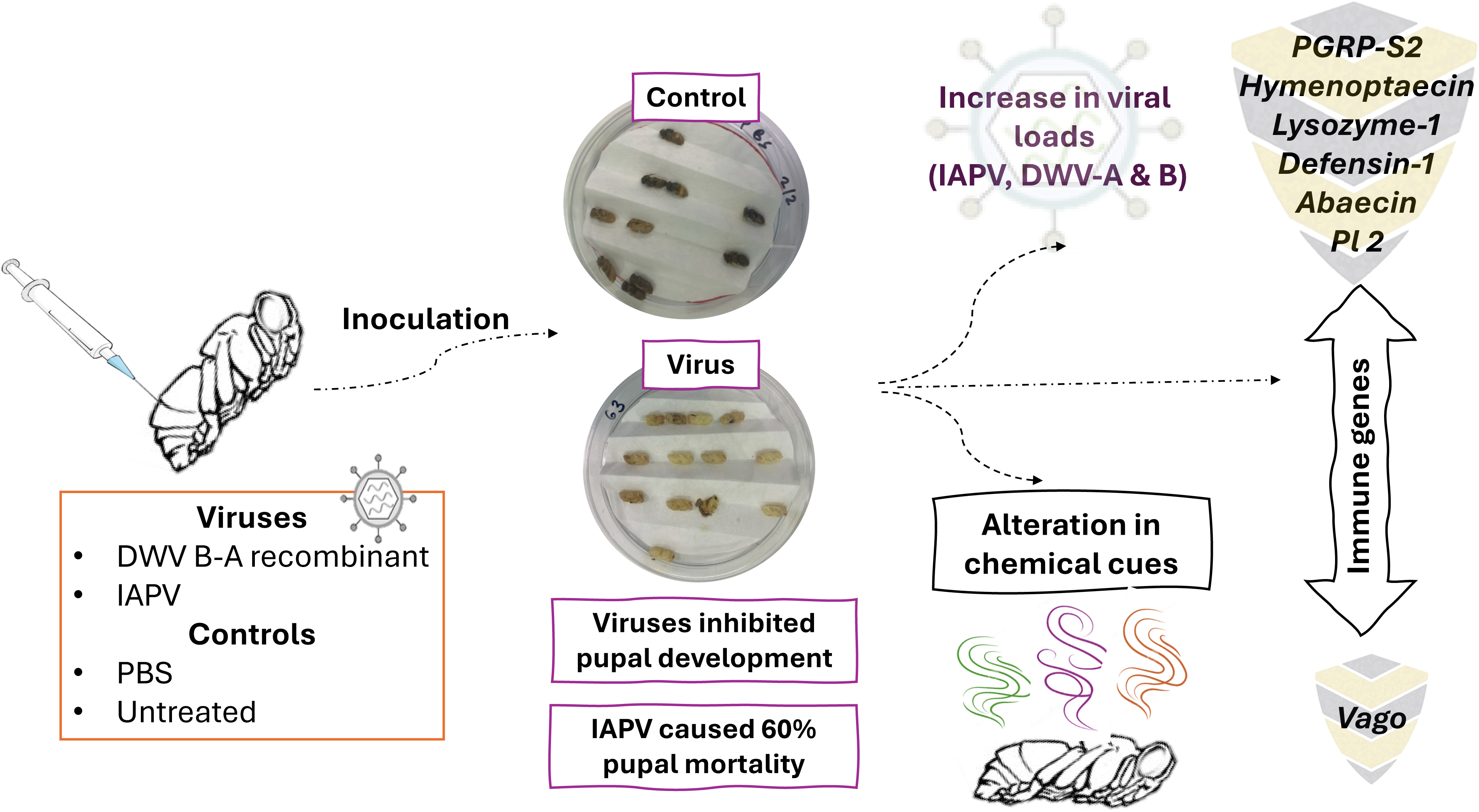

## 1. Introduction

In the last decades, the health of honey bees (*Apis mellifera*) has been a topic of concern (1–3). The combination of the parasitic mite *Varroa destructor* (Varroa) and the viruses it vectors is major driver of honey bee mortality (4–6). Varroa feeds exclusively on honey bees, causing weight loss, weakening of bees’ immune system, and transmitting at least eight pathogenic viruses (7–10). These include mainly the deformed wing virus master variants (DWV-A & DWV-B), acute bee paralysis virus (ABPV), and its variant Israeli acute paralysis virus (IAPV) (2,9). So far, there is no effect treatment against the viruses killing honey bee colonies. A sustainable strategy to reduce disease propagation within the colonies, is to utilize honey bees’ innate defense mechanisms: social and individual immunity (11–13). Hygienic behavior is a well-known form of social immunity that is known to be effective against Varroa mites and a few pathogens (12,14,15). This behavior involves detection of infected or infested brood by adult hygienic bees and their subsequent removal from the colony. Previous studies have indicated that hygienic bees identify diseased brood through olfactory cues (16,17). A prerequisite for effective hygienic behaviour is the production of specific cues by infected pupae followed by olfactory sensitivity and efficient processing of this information in workers (18–20).

So far, some specific changes in volatiles emitted from live brood infected with chalkbrood disease (caused by *Ascosphaera apis*), Varroa mites alone or with DWV infections, and dead brood have been identified (20–24). Brood infected with chalkbrood disease releases three specific volatiles: phenethyl acetate, 2-phenylethanol, and benzyl alcohol that were absent in volatiles of healthy brood (23). Volatiles from Varroa-infested brood were characterized by increased levels of some hydrocarbons: Salvy et al. (2001) (25) found that Varroa-parasitized pupae emitted higher quantities of long-chain unsaturated hydrocarbons: 10-C33:1 (tritriacontene) and 10 + 9-C31:1 (hentriacontene). Wagoner et al. (2019) (24) confirmed increased production of tritriacontene and hentriacontene by Varroa-infested brood. On the other hand, Nazzi et al. (2004) (22) found an increase in two short chain alkenes C15:1 (pentadecene) and C17:1 (heptadecene), while ethyl hexanoate and α-pinene were reported by Liendo et al. (2021) (26) and recently Noël et al. (2025) (27) in Varroa-infested pupae. Moreover, volatiles from brood parasitised by Varroa and infected with DWV have been found to emit higher proportions of 2- and 3-methylbutanoic acid and tritriacontane (24,28). Many of the above-mentioned volatiles were shown to induce hygienic behaviour. However, to the best of our knowledge, there is no evidence to date on whether natural viral infections or virus inoculation, in the absence of Varroa infestation, changes the volatile profile of the brood or induces hygienic behavior.

At the individual level, the immune response of honey bees is well studied. Cellular responses include phagocytosis, melanization, encapsulation mediated by haemocytes, and induction of RNA interference (RNAi) pathway that is triggered by dsRNA generated during the replication of RNA viruses (29). Humoral responses produce antimicrobial peptides (AMPs) by immune gene cascade pathways; e.g., Toll and Imd (Immune Deficiency) involving NF-kB (Nuclear Factor kB) homologues, JAK/STAT (Janus Kinase/Signal Transducer and Activator of Transcription), JNK (c-Jun N-terminal kinase), and MAPK (Mitogen-Activated Protein Kinases) (30–32). In general, the immune response varies across the developmental stages of honey bees (33), and in particular was found to be weaker in brood than in adults, in both workers and drones (34–36). Immune challenges such as Varroa parasitism, and bacterial or viral inoculation were reported to upregulate the expression of immune genes, in both adults and brood. The AMP’s *lysozyme* and *hymenoptaecin* increased in fifth instar honey-bee larvae challenged with *Paenibacillus larvae* (37). *Vago* from the RNAi pathway and *defensin-1* from the Toll pathway increased in DWV-injected pupae (38,39). A higher expression of *defensin-2*, *hymenoptaecin* and *relish* from the Toll and Imd pathways was found in Varroa-infected pupae (40). In contrast, downregulation of genes from the beginning of the Toll pathway cascade such as peptidoglycan recognition protein S2 (*PGRP-S2*) and *spatzle* were detected in pupae infected with Varroa and DWV (39,40). Moreover, immunosuppression of genes from the Toll pathway, under the control of Nf-kB, was found in 5th instar larvae infected with DWV (41). Could it be that downregulation of pupae’s immune response, allows viral replication, that in turn leads to changes in chemical cues, thus promoting pupae ‘altruistic suicide’ by hygienic response for timely removal of infection from the colony? It is also possible, that upregulation of immune response inhibiting viral replication thus could lead to asymptomatic infection, making the virus transmission more effective?

In Israel, a common DWV variant is a recombinant between the DWV-B capsid and DWV-A non-structural genomic regions (42). The recombinant DWV B-A virus was identified in the heads of recently emerged symptomatic bees. Moreover, this alternative DWV genomic arrangement was also observed in the U.K. (43) and more recently in the U.S. in a 2021 survey (44). In this study, we tested the impact of inoculation with DWV B-A recombinant compared to that of an IAPV-enriched inoculum on the volatile profile and immune response of pupae from high-hygienic colony, without Varroa infestation.

## 2. Results

### 2.1 The composition of the circulating virus in the experimental colony

We were only able to assemble contigs but incomplete genomes from the experimental colony that was assigned to DWV and Lake Sinai virus (LSV) (Table. 1S). This indicated that both DWV and LSV were circulating in the experimental colony at the time we collected the pupae for our inoculation experiments. No contigs for sacbrood virus (SBV) nor IAPV could be assembled.

### 2.2. The composition of the virus inocula

Both RT-qPCR (Fig 1A) and ONT sequencing analysis (Table 1.) of IAPV and DWV-enriched inocula revealed that they were not pure. According to RT-qPCR, DWV-A and IAPV were found in both inocula. Moreover, we know that the RT-qPCR targets known viruses but only a short genomic region of the targeted virus, whereas ONT allows detection of full genomes and recombinants (e.g. (45)). Consequently, ONT sequencing analysis found that the DWV B-A recombinant was the main virus (82%) and the only complete virus in the DWV-enriched inoculum (Table 1.). ONT sequencing also confirmed the RT-qPCR result indicating the presence of short contaminating genomic regions of other organisms and viruses, albeit in low abundance. The IAPV-enriched inoculum contained two draft/complete genomes of the AKI complex, namely IAPV and ABPV, in addition to the DWV B-A recombinant (Table 1.).

**Fig. 1.**
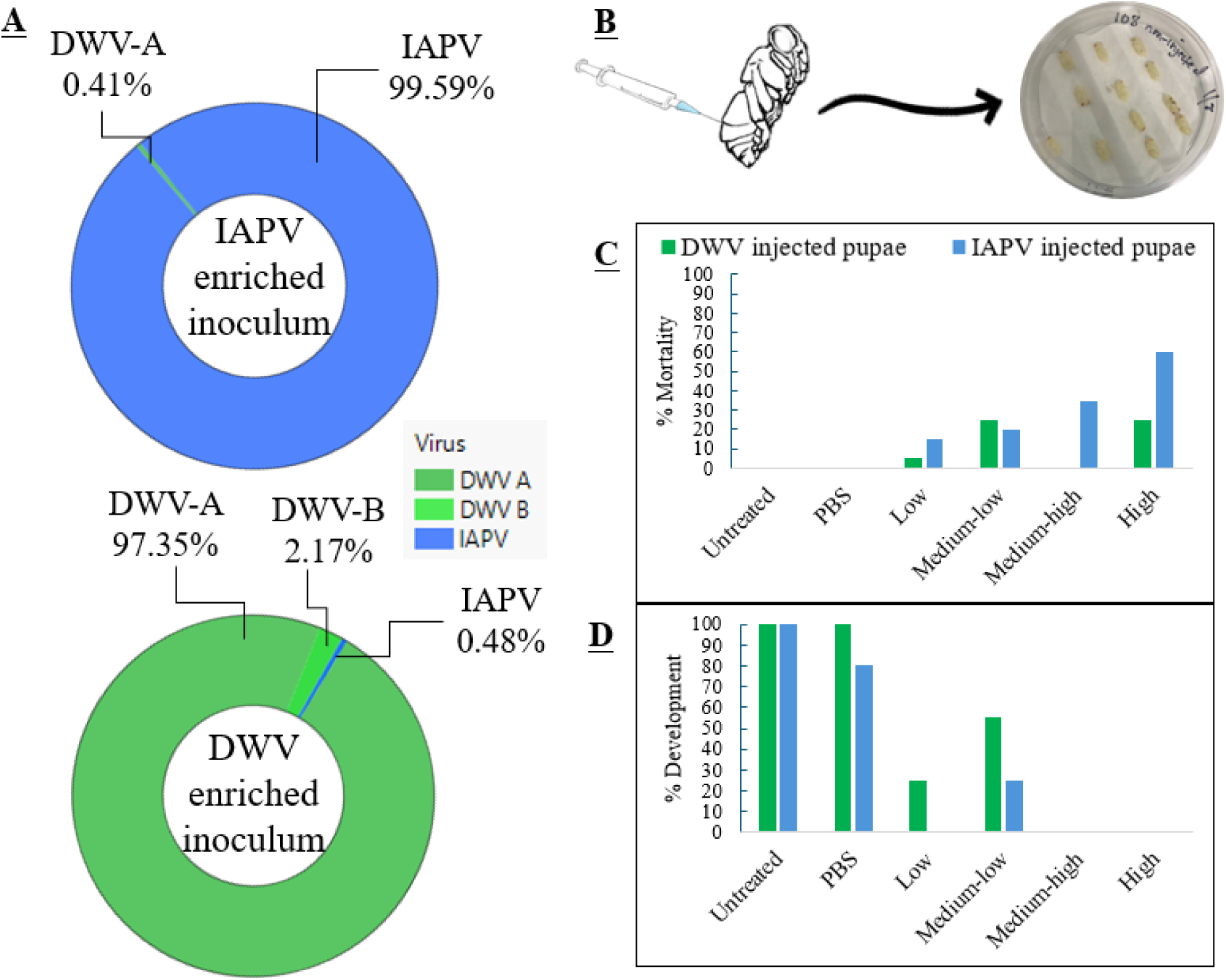
**A.** Proportion of IAPV, DWV-A, and DWV-B in IAPV and DWV-enriched inocula measured by RT-qPCR that targets the RdRp gene (e.g. (46)). **B.** Pupae inoculation: white-eyed pupae were injected with 1µl of either IAPV or DWV-enriched inoculum at four concentrations per inoculum. Injection with 1µl of PBS and untreated pupae were used as negative control treatments. **C.** Pupal mortality percentage 5-days post-treatment, for each treatment. Green bars represent the results of the DWV-injected pupae, and blue bars represent the results of the IAPV-injected pupae. **D.** Percentage of pupae development to the last developmental stage, 5-days post-treatment, in each treatment. Green bars represent the results of the DWV-injected pupae, and blue bars represent the results of the IAPV-injected pupae.

**Table 1.**
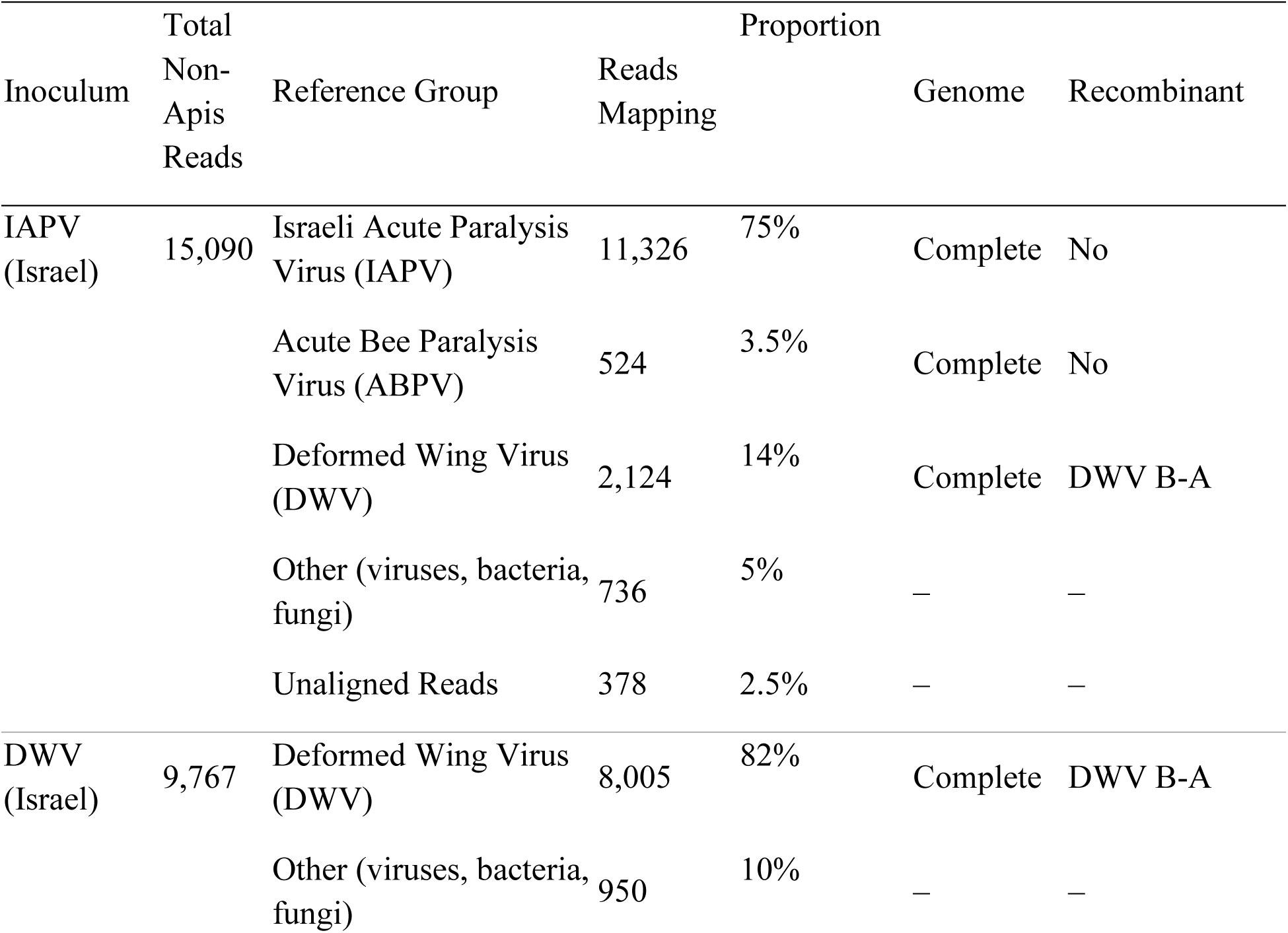
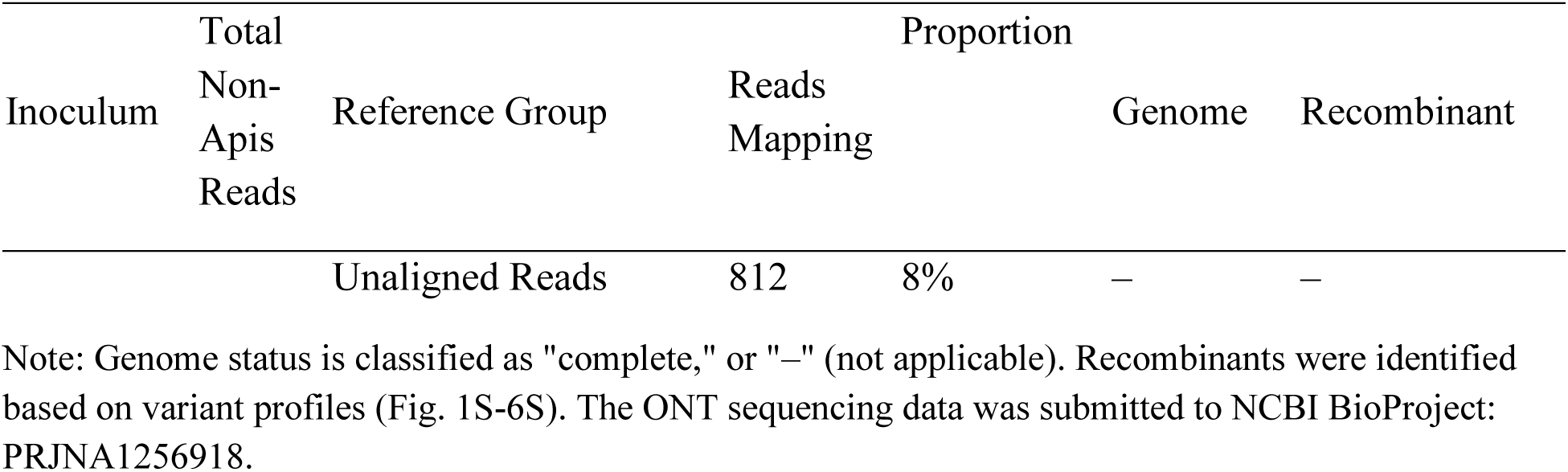
Mapping and assembly of ONT sequencing reads from IAPV and DWV inocula to viral and host genomes, with genome completeness and recombinant detection.

Both RT-qPCR (Fig. 1A) and ONT sequencing (Table 1.) confirmed that the highest titer in the IAPV-enriched inoculum was of IAPV (99.56% of total virus based on RT-qPCR or 75% of the ONT reads assembled to create a complete genome). The highest titer in the DWV-enriched inoculum was DWV with the DWV-A replicase gene (97.35% of total viruses or 82% of ONT reads mapped to the DWV B-A recombinant complete genome), while genomic copies of DWV-B were present only in the DWV-enriched inoculum based on the RT-qPCR analysis (Fig. 1A) but was a minor proportion of the 10% ONT “other” reads that could not be assembled into a complete genome (Table 1).

### 2.3 Effect of viral infection on pupae mortality and development

No mortality occurred in the control groups in both trials (untreated and PBS mock-injected) (Fig. 1C). IAPV-injected pupae had a dose-dependent and higher mortality rate than DWV-injected pupae. The viral infection also affected pupae’s development (Fig. 1D). In the DWV trial, mock injected pupae developed as well as the untreated, reaching 100% relative development, whereas low and medium-low DWV-injected pupae reached only 25% and 55% relative development, respectively. In the higher DWV concentrations, development was inhibited in all the pupae. In the IAPV-trial, the development of mock injected was delayed in 20% of pupae relative to untreated control, and only 25% of the pupae from the medium-low treatment developed to the stage of untreated control, while development was inhibited in all the other IAPV treatments.

### 2.4 Effect of viral infection on DWV and IAPV loads by RT-qPCR

The experimental colony was reported as “low virus” according to the ONT results (section 2.1). The controls in both DWV (Fig 2 A & C) and IAPV (Fig 2 B & D) trials, namely the untreated and mock PBS-injected pupae, contained similar amounts of DWV-A with the only exception being the absence of detection in the mock-injected DWV controls (Fig 2 A). The low levels of DWV-A circulating in the colony were anticipated based on the ONT results, however, the IAPV loads indicate that the ONT sequencing was carried out at a sequencing depth that did not detect the presence of a viable source of IAPV. The loads in control pupae were significantly lower than those of the virus-injected pupae (ANOVA: F (_5,72_) =18.6, p<.0001; Table 2).

**Fig. 2.**
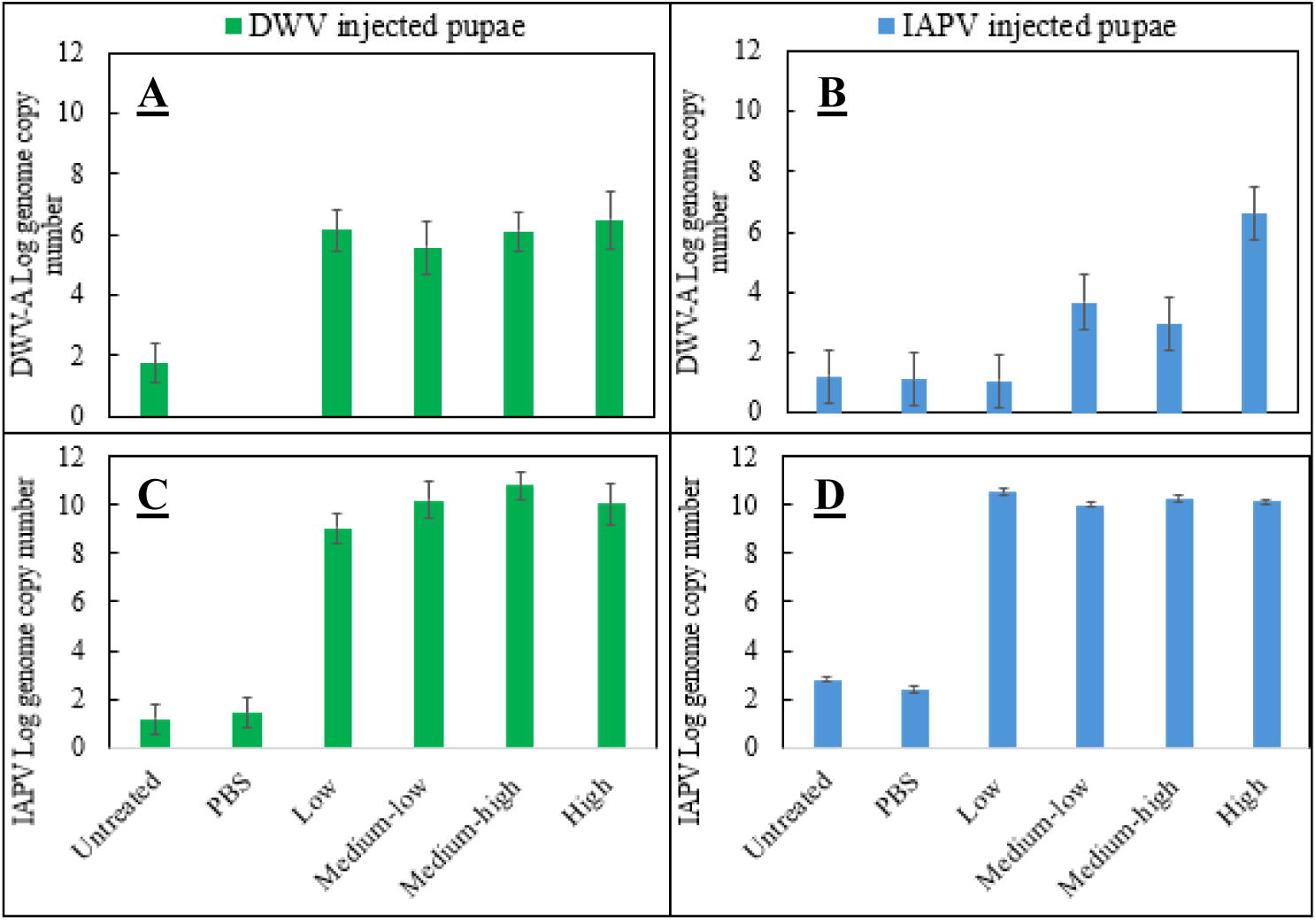
Each graph represents the log-transformed loads of viruses in each of the six treatments (two controls – untreated and PBS mock injected; four viral doses), five days post-treatment. Green bars represent the results of the DWV-injected pupae, and blue bars represent the results of the IAPV-injected pupae. **A.** Loads of DWV-A in DWV recombinant injected pupae; **B**. Loads of DWV-A in IAPV-injected pupae; **C.** Loads of IAPV in DWV recombinant injected pupae; and **D**. Loads of IAPV in IAPV-injected pupae. The bars represent means ± SE of five pupae for each treatment/virus.

**Table 2.**
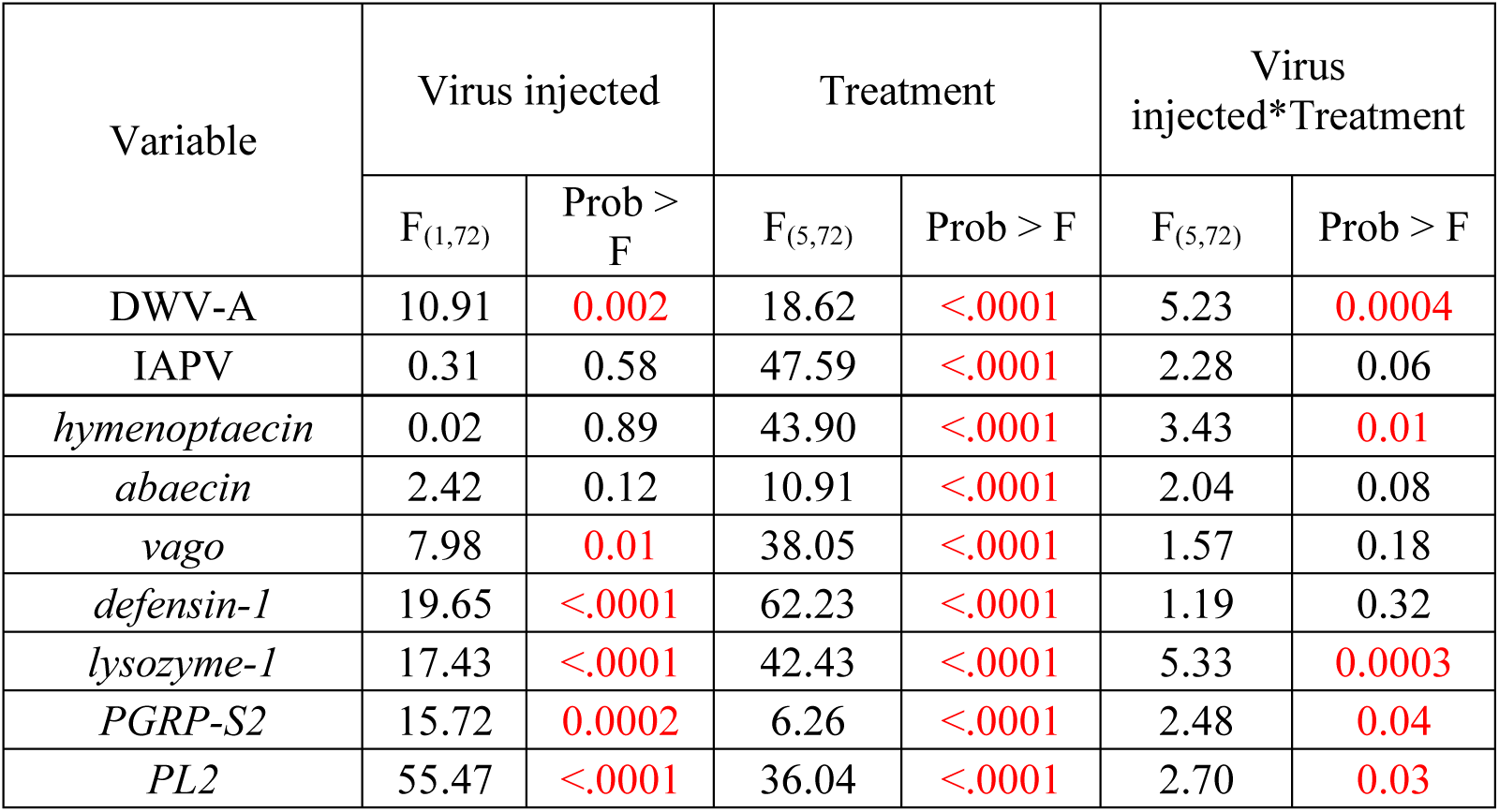
Two-way ANOVA on the ranks of viral loads and immune gene expression and the different effects on the variables: Virus injected (DWV recombinant or IAPV) and Treatment (untreated, PBS-injected, and four viral doses: low, medium-low, medium-high and high). Viral loads and the expression of immune genes were Log transformations prior to analysis.

Five days post-injection, although pupae were injected with different doses of each inoculum, the loads of DWV-A increased in most groups of virus-injected pupae but were higher in DWV B-A recombinant injected pupae than in IAPV-injected pupae (ANOVA: F _(1, 72)_ =10.9, p=0.005; Table 2; Fig. 2 A & B respectively). Moreover, no dose-dependent effect was found in the DWV-A loads in pupae injected with different concentrations of the DWV inoculum (Fig 2 A). Interestingly, DWV-A loads in the IAPV-injected pupae changed significantly with dose (Fig. 2B). Moreover, a significant difference was found between treatment “high” virus to all other doses (Anova, F _(5,36)_ =13.1, p<0.0001 followed by Tukey-Kramer; Fig. 2B). In addition, IAPV loads were similar in both IAPV and DWV-injected pupae (ANOVA: F _(1, 72)_ =0.3, p=0.58; Table 2. & Fig. 2 C&D), irrespective of dose.

### 2.5 Effect of viral infection on pupae chemical cues

GC analysis of the pupal cuticular extracts revealed 157 peaks, whereas GCMS analysis revealed that many of these peaks contained more than one compound not clearly separated (Fig. 7S). Out of 157 peaks detected by GC, 92 were tentatively identified based on their MS spectra. In the extracts we identified members of four classes of hydrocarbons, including n-alkanes, methyl-alkanes, dimethyl alkanes and alkenes with chain-lengths of the carbon ranging from n-C_14_ to n-C_33_, as well as fatty acids, esters as well as a few unknown compounds (Table 3S). It is important to note that analysis of pooled samples from different treatments by GCMS revealed that long-chain fatty acids were not detected in untreated and PBS-injected pupae. The most abundant n-alkane for all treatments was heptacosane (n-C_27_), where is long-chain fatty acids were found almost exclusively in the extracts of virus-injected pupae. ANOVA analysis followed by Bonferroni correction of individual GC profiles revealed that the intensity and abundance of the volatiles varied among treatments. In particularly 20 were found significantly different by treatment (Table 3S). Among these, relative quantity of n-C_21_ was significantly higher in untreated and PBS-injected pupae (ANOVA, F_(9, 49)_=8.54, p<0.0001, Tukey p< 0.05), whereas the relative quantity of octadecanoic acid was higher in injected pupae vs. control (ANOVA, F_(9, 49)_=8.07, p<0.0001, Tukey p<0.05). Interestingly, the relative quantity of unidentified peak 67 was highest in DWV high group and significantly different from all IAPV-injected pupae (ANOVA, F _(9, 49)_ =7.19, p<0.0001, Tukey p< 0.05). Principal component analysis on relative quantity of these 20 peaks discriminated between virus-injected (IAPV or DWV) pupae and the control pupae (untreated and mock PBS-injected) (Fig. 3). Profiles of the untreated and PBS-injected pupae clustered together, indicating a similar volatile profile, while the volatile profiles of IAPV and DWV-injected pupae clustered somewhat separately (Fig. 3). Especially high separation was between the profile of pupae injected with high concentration of DWV and the other groups.

**Fig. 3.**
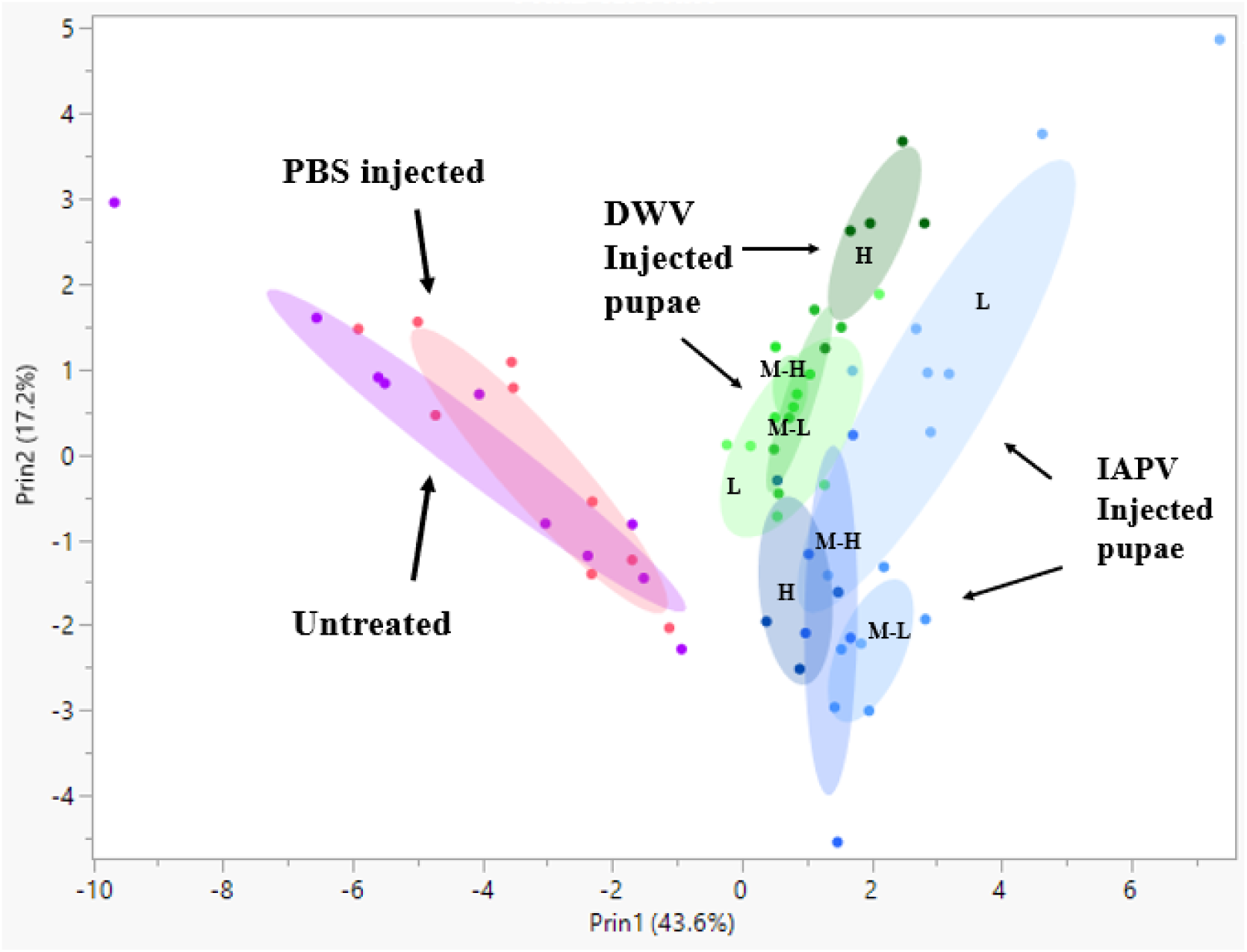
Principal component analysis on 20 volatiles secreted by pupae, five days post-treatment. The analysis was conducted on peaks whose relative quantity was significantly different by ANOVA after Bonferroni correction p<0.0003. Different colors indicate treatments as labelled.

### 2.6 Effect of viral infection on pupae expression of immune genes

The expression of all the tested immune genes: *vago, hymenoptaecin, abaecin, defensin-1, lysozyme-1, PGRP-S2,* and *Pl2* was affected by the treatments (Table. 2). In particular, virus-injected pupae had significantly different immune gene expressions compared to both controls, PBS-injected and untreated pupae in both trials (Table. 3). Interestingly, *vago* was the only immune gene that was downregulated in the virus-injected pupae of all groups (Fig. 4A), while *hymenoptaecin* was upregulated in most virus-injected groups (Fig. 4B). All the other immune genes had higher expression following virus injection vs. control treatments (Fig. 4C-G). The expressions of *vago*, *defensin-1*, *lysozyme-1*, *PGRPS2*, and *Pl2* were higher in the DWV-injected pupae than in the IAPV-injected pupae (Fig.4A, C, E, F, and G;Table 2.); whereas the expressions of *hymenoptaecin* and *abaecin* were not affected by the type of virus injected (Fig.4 B and D;Table 2.) and showed a similar response to both injected viruses. *Pl2*, *lysozyme-1,* and *PGRP-S2* were significantly affected by the interactions. The expression of *Pl2* was significantly upregulated in the medium-low DWV-injected pupae than in the medium-low IAPV-injected pupae (Fig. 4C;Table 2.). The expression of *Lysozyme-1* was significantly downregulated in the medium-high IAPV-injected pupae than the medium-high DWV-injected pupae (Fig. 4F; Table 2.).

**Fig. 4.**
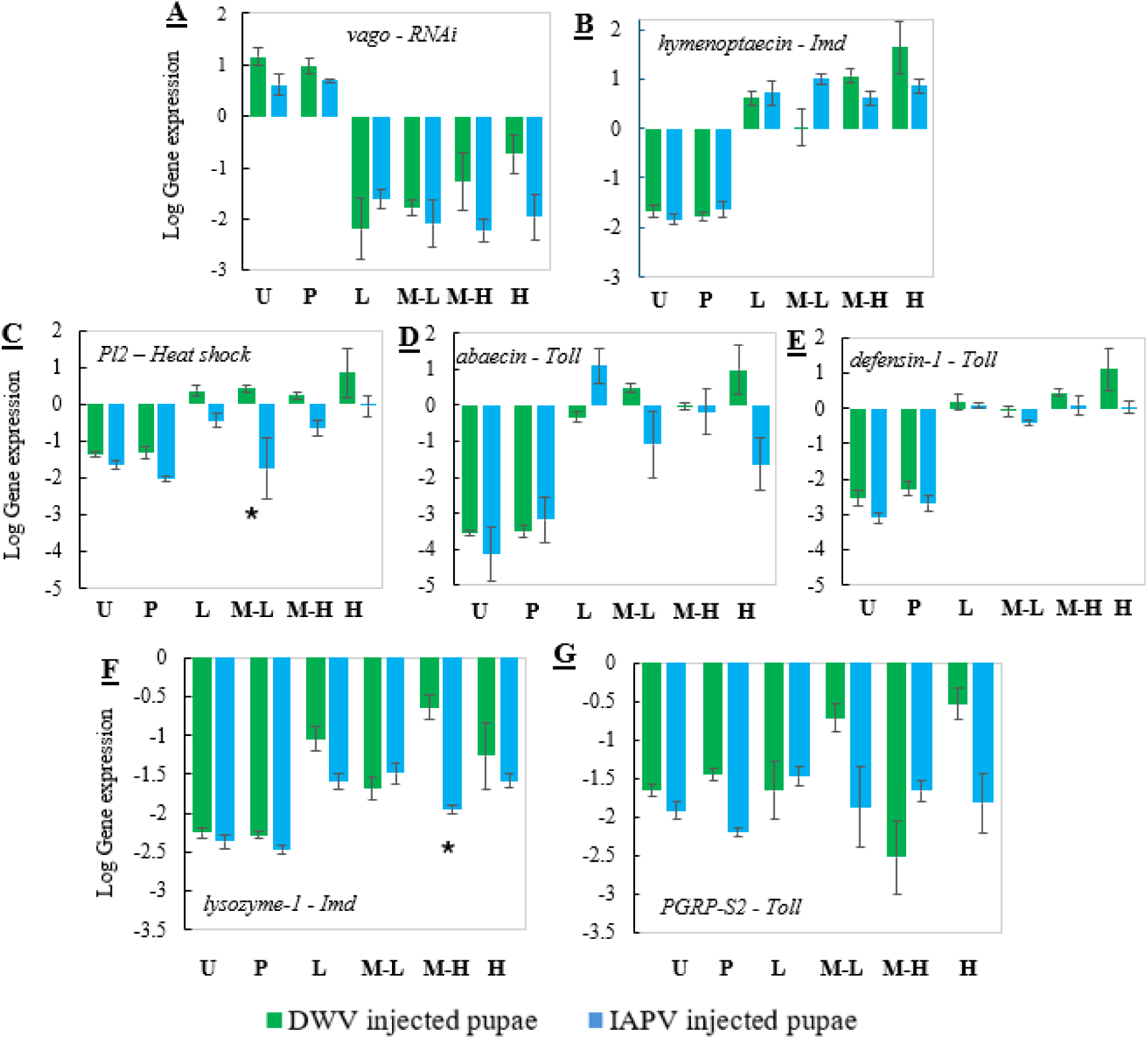
Each graph represents the log-transformed expression of the immune gene, by treatment: U-untreated, P-PBS-injected, and four viral doses: L-low, M-L-medium-low, M-H-medium-high and H-high. **A.** *vago*, **B.** *hymenoptaecin*, **C.** *Pl2*, **D.** *abaecin*, **E.** *defensin-1*, **F.** *lysozyme-1*, **G.** *PGRP-S2*. DWV-injected pupae are represented by green bars and IAPV-injected pupae by blue bars. The bars represent means ± SE of five pupae for each treatment/virus and *-represents significant differences between the trials, injection with either IAPV or DWV-inoculum, (Two-way Anova, p<0.05).

**Fig. 5.**
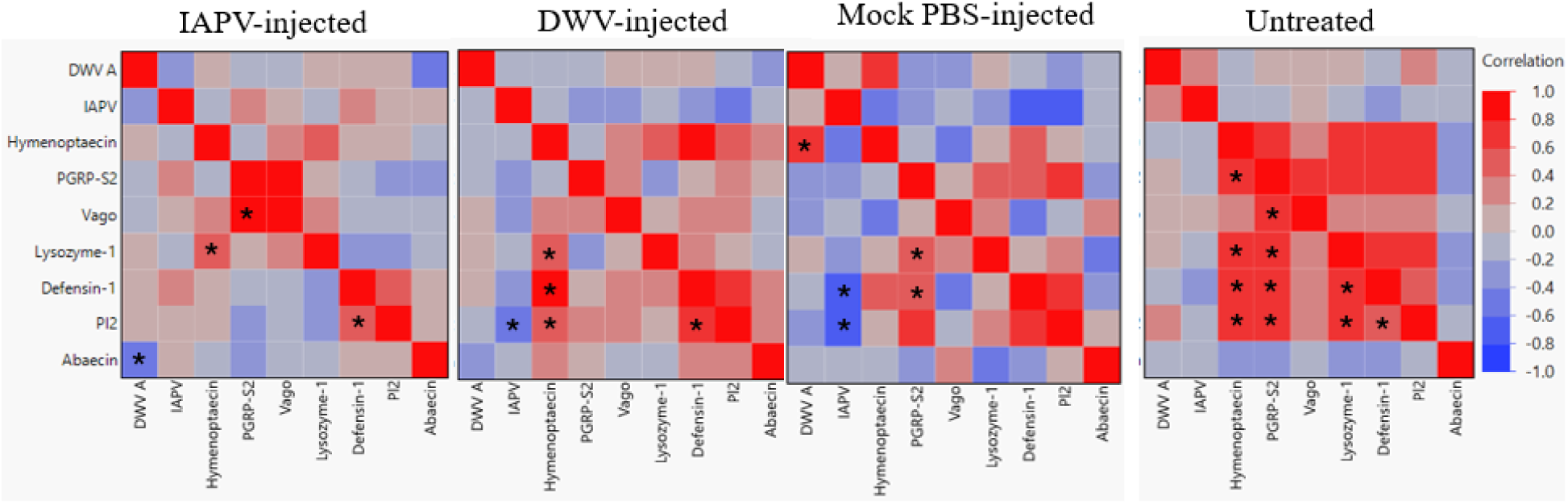
Heat map correlation between viral loads and expression of immune genes in pupae either injected with IAPV, DWV and PBS or left untreated: IAPV-injected pupae all pooled together (on the left), DWV recombinant injected pupae all pooled together (in the middle-left), mock PBS-injected pupae pooled together (in the middle-right) and untreated pupae all pooled (on the right). The viral loads and expression of immune genes were log-transformed.

### 2.7 The interactions between viral loads and expression of immune genes in virus-infected pupae (IAPV- or DWV-infected) vs. the control pupa (untreated and PBS-injected pupae)

Correlation analysis between viral loads and expression of immune genes revealed differences between virus-infected pupae (IAPV- or DWV-infected) vs. the control groups, PBS-injected and untreated pupae. The untreated pupae had the highest number of significant correlations between the immune genes, and all were positively correlated. *Hymenoptaecin* and *PGRP-S2* had significant positive correlations with each other and with *lysozyme-1*, *defensin-1* and *Pl2*. *PGRP-S2* also had a positive correlation with *vago*. *Lysozyme-1* positively correlated with *Pl2* and *defensin-1*. *Pl2* and *defensin-1* were also positively correlated. Injection with PBS changed dramatically the interactions and left only two correlations between the immune genes: positive correlation between *PGRP-S2* with *lysozyme-1* and *defensin-1*. Whereas viral loads correlated with some immune genes. IAPV loads negatively correlated with *Pl2* and *defensin-1*, and DWV loads positively correlated with *hymenoptaecin*. DWV-infected pupae had some similar significant correlations between the immune genes as the untreated pupae; *hymenoptaecin* positively correlated with *lysozyme-1*, *defensin-1* and *Pl2*. *Pl2* and *defensin-1* were also positively correlated, and *Pl2* and IAPV loads negatively correlated. IAPV-infected pupae had the lowest number of significant correlations. Positive correlations were found between *vago* and *PGRP-S2*, between *hymenoptaecin* and *lysozyme-1*, and between *Pl2* and *defensin-1*. DWV-A loads were negatively correlated with *abaecin*.

## 3. Discussion

The global decline of honey bee populations in 2006-2007 (1) has been well documented and the more recent January 2023 collapse in commercial bee operations in the U.S. is an ongoing concern (47). It was reported that over 60% of commercial beekeeping colonies had been lost since the prior summer of 2024, representing 1.7 million fewer colonies available for pollination services. The role of honey bees as pollinators is vital to both the world’s environment and economy. As it was in the 2006-2007 honey bee global declines (48), viruses that are often vectored by Varroa also played a key role in colony losses in 2023 (47). They reported that high levels of DWV-A and B and another member of the AKI complex, acute bee paralysis (ABPV), dominated the virus community as tested by RT-qPCR. No sequencing was carried out in that study, however, viral recombinants between DWV A & B could be involved as Hesketh-Best et al., (2024) (44) reported the dominance of recombinant DWV genomes in a national U.S. honey bee and varroa mite survey carried out in 2021. A similar observation of increasing prevalence of recombinants between DWV A & B was reported in Europe (49), but as pointed out by Paxton et al. (2022) (50), the role of the recombinants remains largely ambiguous. In Israel, the DWV B-A recombinant was first detected by Zioni et al. (2011) (42) in the heads of recently emerged symptomatic bees. This DWV B-A recombinant has the combined features of the DWV-B capsid genes and DWV-A non-structural genomic regions (42). Honey bees in Israel are also often infected with the AKI complex, to a greater extent with IAPV rather than ABPV (51,52). While the many surveys implicate DWV A & B, their recombinants and the co-infection with AKI complex viruses with honey bee losses, we have yet to experimentally observe the impact of such infections on honey bees brood in the absence of Varroa. And whether these infections elicit volatile chemical profiles and immune response in pupae that could be a benefit in a high-hygienic colony.

Our results show that viral inoculation in the absence of Varroa infestation had a dramatic effect on pupal survival and development. Both IAPV and DWV B-A recombinant enriched inoculum injections caused delays in development and increased mortality. The IAPV inoculum had the most profound impact on survival and development in a dose-dependent manner. Most of the IAPV-infected pupae in the low treatment died and none developed, while injection with mock PBS alone had no negative effects. These data indicate that the IAPV inoculum was more virulent than the DWV B-A inoculum. In fact, both inocula were not pure. According to ONT analysis, IAPV-enriched inoculum contained two draft/complete genomes of the AKI complex, namely IAPV and ABPV, in addition to the DWV B-A recombinant, while the DWV inoculum contained complete genomes only of DWV B-A recombinant. However, according to RT-qPCR, DWV-A (detecting the target replicase region of the DWV B-A recombinant) and IAPV were found in both inocula, albeit with different prevalences. As the RT-qPCR design only targets a short replicase genomic region of DWV, ONT sequencing proved more insightful as it allows detection of complete or near complete genomes and possible recombinants, as we subsequently observed in our inocula. This enhanced clarity was previously eloquently highlighted by Nikulin et al. (2024) (45). Future surveillance and indeed laboratory-based manipulation studies should consider including some aspects of whole genome sequencing. It not only provides valuable information about the virus genomes, their mutations and genomic rearrangements, but any co-infections that might be co-occurring.

Our RT-qPCR analysis revealed that injection of either inoculum resulted in increased loads of both DWV-A and IAPV. The identity of “DWV-A” in pupae injected with IAPV is quite clear. It is most likely the DWV B-A recombinant known to be present in the IAPV inoculum. However, we cannot exclude the possibility of another endogenous DWV-A variant co-circulating in pupae at the time of pupae collection, as was observed in the control groups. Here-on-in we will refer to the DWV-A RT-qPCR positive result a proxy for the detection of the DWV B-A recombinant as delivered in the inocula since a dose-dependent amplification of DWV B-A by IAPV inoculum supports the idea that its main source of the DWV infection. The source of the IAPV in DWV B-A injected pupae is less clear. Surprisingly, IAPV loads reached maximum levels, irrespective of DWV dose applied, reaching levels at day five equivalent to the IAPV, levels as observed in the IAPV-injected pupae. The IAPV likely originated in the pupae’s endogenous covert infections, as we found low loads of IAPV in the untreated pupae. However, the injection itself did not raise the IAPV internal viral loads, as the IAPV loads in mock PBS-injected pupae remained low and similar to the untreated pupae. An additional possibility is that the source of IAPV was in the inoculum itself, as found by RT-qPCR. A similar scenario was reported by Carrillo-Tripp et al. (2016) (53) when emerging bees and AmE-711 cell line (derived from honey bee embryos) were infected with enriched sacbrood virus (SBV) inoculum. The infected bees and the cell line resulted in IAPV infection, although the percentage of IAPV in the SBV inoculum was less than 1%. However, the fact that our ONT analysis did not detect the full sequence of IAPV questions this possibility, although it could be a result of the sequence depth applied in our sequencing effort. For example, transcriptome studies which follows a similar workflow as detecting RNA viruses noted that genomic variants can be missed even when 30-50X genomic depth is achieved (54). Disregarding the origin of IAPV, the question is how the levels of IAPV in DWV B-A injected pupae reached similar loads to IAPV-injected pupae? It could be a result of a combination of two factors, the immune suppression by the DWV injection (41,55), followed by the high replication speed of the latent IAPV (56–58). In fact, we found that the expression of immune genes in virus-infected pupae five days post-injection changed in a quite complicated manner.

We tested the surviving pupae for the expression of immune genes from several pathways: the humoral response (*PGRP-S2*, *hymenoptaecin, lysozyme-1, defensin-1, abaecin*), heat shock response (*Pl2*), and RNAi interference (*vago*), in virus-injected, mock PBS-injected, and untreated pupae. Like in previous reports, all the tested pathways responded to viral infections (39,59,60), and their expression was significantly different from control pupae (mock PBS-injected and untreated). In our experiment, only *vago* from the RNAi pathway was downregulated in both IAPV and DWV B-A inoculated pupae, whereas the expression of the other immune genes increased. *Vago*’s expression in IAPV-injected pupae was lower than that of DWV B-A injected ones. It could indicate the inhibition of RNAi pathway, thus enabling IAPV replication as previously suggested by Chen et al. (2014) (60). An inhibition in expression of immune genes was also reported before (41,59,61,62) and it was considered as an immunosuppression strategy of the virus to evade the host’s defenses as a part of well-known evolutionary arms race between viruses and the immune systems of their hosts. However, all the other tested immune genes were upregulated following viral inoculations but not in the same manner. The most induced transcript was the antimicrobial peptide from the IMD pathway-*hymenoptaecin*. The AMP *lysozyme-1* from the same pathway was also upregulated, though in less clear manner. All the three transcripts from the Toll pathway, *PGRP-S2*, *defensin-1*, and *abaecin* were upregulated following viral infection. Moreover, their expression appeared higher in DWV B-A infected pupae. Also, the heat shock protein, *Pl2*, was upregulated following viral infections of as was previously found (63–66) and again its induction towards DWV injection was higher. Could the stronger immune response in DWV B-A infected pupae be a result of higher loads of both IAPV and DWV recombinant or better reaction of the immune system to DWV recombinant infection, which also resulted in higher survival? It can be speculated that the latter phenomenon could be explained by the longer history (coevolution) between the honey bee and the DWV viruses. It is important to note that the mock injection did not alter the levels of the tested immune genes, indicating that all the recorded responses were not related to mechanical damage by injection or infusion of isotonic solution, but a response to the pathogen.

Significant correlations between the expressions of most tested immune genes in the untreated pupae, but naturally infected by low level of viruses, indicate crosstalk among all the pathways, suggesting that these pathways were acting together in these pupae, keeping the viruses at an asymptomatic state, allowing continuation of pupae development despite infection. Mechanical damage (i.e. mock PBS-injection) and viral infection with DWV B-A or IAPV inocula altered the interactions between these genes but in different manners. In fact, following mechanical damage, positive correlation remained only between three genes from Toll and IMD pathways (*PGRP-S2*, *lysozyme-1* and *defensin-1*), while correlation between immune genes and viral loads occurred. *Hymenoptaecin* became positively correlated with the DWV-A loads, while others and most *Pl2* and *defensin-1*became negatively correlated with loads of IAPV. This situation indicates that the injection had an impact, but the pupae were able to keep the viruses under control, as the viral loads in these pupae remained similar to those of the untreated ones. Once we added viruses by artificial inoculation, the immune system was unable to keep the viruses under control. Following injections of IAPV and DWV B-A recombinant, the interactions between the immune genes and between viruses and immune genes were different. In particular, *Pl2* was negatively correlated with IAPV levels in DWV B-A injected pupae, and *abaecin* was negatively correlated with DWV recombinant loads in IAPV-injected pupae. Since IAPV loads were similar in virus-injected pupae, the biological significance of these interactions is hard to interpret. These findings emphasize the complexity of the honey bee’s immune response to viral infection in the surviving pupae. It could be that the entire response involved additional immune pathways that we did not test in this study; future deep sequencing and gene manipulation experiments will be required to expose the impact of the immune mechanisms on viral infection in honey bee pupae.

Our experiment showed that although some of the infected pupae died, many survived in laboratory conditions. We wondered if these virus-infected pupae could survive in the colony or would be detected by hygienic bees and removed from the colony. It has been reported that hygienic behavior is induced by high and low volatile compounds towards Varroa-infested brood (20,22,26–28). We focused on cues from virus-infected pupae in the absence of any traces of Varroa infestation. Our data indicate that the injection itself did not change the pupae’s chemical cues, whereas viral infection did. These findings do not contradict the fact that the injection per se can induce changes in pupae volatiles shortly after piercing, as we tested the pupae five days after injection. In particular, the relative amount of 20 compounds was different between virus-infected surviving and control pupae (untreated and mock PBS-injected), thus contributing to the “unhealthy” or “unpleasant” pupae smell. Among them, the fatty acids (palmitic (C_16_) and stearic (C_18_) acids) were the most prominent according to the MS analysis, in pupae that were injected with high doses of either IAPV or DWV B-A. According to analysis of pooled extracts additional acids of lower molecular weight were also limited to pupae injected by high loads of viruses. However, we failed to detect these in individual samples probably to their low abundance. Interestingly, ethyl oleate, a known primer honey bee pheromone (67) was highest in virus injected pupae. This finding is surprising and will require a follow up study. The other distinct volatiles were hydrocarbons, such as heneicosane which was more prominent in control pupae. It is worth noting that we did not detect any volatiles previously identified in Varroa-infested brood (20,28). How specific these changes in volatiles to viral infection is hard to tell now. Long-chain fatty acids were not reported in Varroa-infested brood, while particularly oleic acid, was suggested as the death pheromone in honeybees from freeze-killed brood at different developmental stages; moreover, it is apparently detectable by hygienic workers, inducing hygienic behavior (21,68). However, in our experiments, the pupae were not dead, but experienced retarded development. We detected some changes in relative quantities of a few compounds not previously associated with Varroa infestation or hygienic behavior but not in high and low volatile compounds detected in virus-infected pupae reported by others. The inability to detect high volatiles could be explained by the differences in the methodology used, volatile collection (21,26–28) and solvent extraction. However, the fact that we did not detect differences in relative quantities of unsaturated alkenes n-C31:1 and n-C33:1 reported by (24,25) in Varroa and DWV could be attributed to differences in cues emitted by virus and Varroa-infected pupae but not methodology. An alternative explanation could be that the chemical cues to DWV-A or B is different to those elicited by our DWV B-A recombinant. This remains an open question that needs to be addressed in future experiments.

Since we excluded dead pupae from the analysis, we hypothesize that the volatiles detected here were emitted by the pupae surviving the viral infection and thus may serve as a cue or a signal for the uncapping and removal by hygienic workers. Whether the hygienic workers use these cues in the absence of cues from the Varroa mite as a signal to activate hygienic behavior remains to be tested. Would the hygienic response curtail or increase virus transmission throughout the colony? With regards to the DWV B-A recombinant, are the chemical cue so different that the bees don’t react to it as if it would to wild-type variants or acute infections as observed for IAPV? Could this explain why the DWV recombinants becoming more prevalent in honey bee colonies in the U.S. and Europe (44,49)? These questions will require further studies.

In conclusion, our findings demonstrate a complicated immune response in honey bee pupae that survived viral infection for five days without Varroa infestation. The response was composed of both induction and inhibition of immune genes and changes in the emission of chemical cues. Whether virus-induced changes in volatiles activate workers’ hygienic behavior within the colony is being tested? In this research, we focused on two key groups of viruses from DWV and AKI complexes. However, colonies are typically infected by multiple types of viruses, other pathogens and affected by other stress factors and their interactions. The immune responses will change, adapt and react accordingly. Deep whole genome sequencing along with gene manipulation is needed to resolve these interactions. Developing strategies to enhance honey bee resilience, particularly through selective breeding for hygienic behavior traits such as improved immune function, is crucial toward sustaining healthy and productive colonies.

## 4. Materials and methods

### 4.1 Colony description

The research was conducted in an experimental apiary at the Volcani Center, Agricultural Research Organization (ARO), Israel. The experimental colony was treated to control Varroa using Amitraz-loaded strips (Apiraz, Luxeburg, Israel) as advised by the extension services. Winter treatment was terminated in mid-February. At the time of the experiments, the colony’s Varroa infestation was assessed by counting the natural mite fall once a week on a sticky board placed on the bottom board of the colony, and the results in the experimental hives were relatively low, on average 5 mites per day. DWV injection experiment was conducted at the end of April, and IAPV injection experiment in the middle of May, in 2023.

The source population of honey bees was a local mixed population mainly composed of *Apis mellifera ligustica*, which has undergone a bidirectional selection program for high (H) and low (L) hygienic responses since 2012 (69) based on the pin-killed brood (PKB) assay, as in Erez et al. (2022) (51). Hygienic colonies were selected based on the results of two sequential 24-hour tests. Colonies were considered hygienic if out of 100 capped and pierced pupae, 98% were uncapped and 95% completely removed.

### 4.2 Inoculum preparation

IAPV-enriched inoculum was prepared as described by de Miranda et al. (2013) (70) with a slightly modified virus purification. In brief, approximately 300 white-eyed pupae were injected into the 2^nd^ and 3^rd^ integuments of the abdomen, with 1µl of IAPV inoculum, diluted with PBS to 10^5^ viral particles/µl. After incubation at 34 °C, 70% RH for 3 days, the pupae were macerated in the extraction buffer (sodium phosphate 0.01 M, pH 7 containing 0.02% sodium diethyldithiocarbamate). The tissue debris was mixed with chloroform and clarified by three centrifugation steps at 8000g at 4 °C for 15 min in Hettich Universal 32R centrifuge (Hettich Lab Technology, Germany). The supernatants were overlaid on a solution of 30% sucrose in phosphate-buffered saline pH 7.5 (PBS) and subjected to 4 hours of ultracentrifugation at 100,000 g at 4 °C in a Sorvall Discovery 90SE ultracentrifuge using a TY35 rotor (Hitachi). The viral pellet was re-suspended in 30% sucrose solution in sterile PBS and kept in -80 °C.

DWV-enriched inoculum was prepared as described in Levin et al. (2016) (71) from about 800 Varroa mites, collected from four untreated honey bee colonies, in November and December 2023.

Both inocula were diluted 1:5 in phosphate-buffered saline (PBS), heated at 95 ◦C for 4 min and cooled on ice for 3–5 min before cDNA preparation (72). Inocula cDNA was quantified via RT-qPCR. Additionally, 24µl of each inoculum was combined with 176 µl of 1X PBS, and the RNA was extracted with Viral Nucleic Acid Extraction Kit II (Geneaid). The RNA samples were sequenced using the Oxford Nanopore Technologies (ONT) sequencing platform (section 4.3).

### 4.3 Oxford Nanopore Technologies sequencing and bioinformatics

Sequencing and subsequent genome assembly were carried out as described previously (45,73). Briefly, double stranded cDNA was created using N6 modified primer and a Template Switching Reverse Transcriptase (New England Biolabs, MA, USA) for the synthesis of the 1st strand. A long-read polymerase (PrimeSTAR GXL polymerase, TakaraBio, USA) was used for the synthesis of the 2nd strand. The cDNA was prepared for sequencing using the ONT Rapid Barcoding Sequencing Kits (SQK-RPB004, Oxford Nanopore Technologies, UK) as per the manufacturer’s guidelines. Libraries were pooled on FLO-MIN106 flow cells and run on the GridION. Sequencing performance was monitored and was terminated after 24 h.

Sequencing reads were filtered to a minimum length (≥200 bp) and Q-value (≥9) by MinKNOW v4.3.4. Basecalling and demultiplexing were performed using Guppy v6.4 with high-accuracy model. PoreChop v0.2.4 was used to remove the nanopore barcode adapter sequences. To assemble contigs, the quality-filtered reads were assembled by Canu v2.2 using the following assembly parameters: *-nanopore maxInputCoverage = 2000 corOutCoverage = all corMinCoverage = 0 corMhapSensitivity = high minoverlap = 50 minread = 200 genomesize = 5000*. Contigs with a minimum length of 1 kbp were binned manually with the anvi’o v7.1 interactive interface. The reads were mapped to the contig database using Minimap2 v2.24, and the read recruitment was stored as a BAM file using samtools. The contigs database was populated with additional data, incorporating HMMER results against Virus Orthologous Groups (VOGs; https://vogdb.org/) in addition to the standard anvi’o HMMR profiles, NCBI COGs and KEGG KOfam database. Contig taxonomy was predicted by running Kraken2 v2.1.2 using the non-redundant NCBI database on the gene calls. Finally, merged profiles were clustered with the automatic binning algorithm CONCOCT, and the anvi’o profile was visualized for manual binning. Binning was guided by sequence composition similarity (visualized as a dendrogram in the Anvi’o interface), and the presence of viral HMM hits to the VOG database. The ONT sequencing data was submitted to NCBI BioProject: PRJNA1256918.

### 4.4 Pupae injected experiment

Total RNA from bee samples from 40 high and low-hygienic colonies was sequenced using the Oxford Nanopore Technologies (ONT) sequencing platform (section 4.3). The experimental colony (#108) was selected from the tested colonies based on its high hygienic behavioral response with known low virus abundance according to the ONT analysis (Table. 1S). White-eyed pupae were collected from the same hygienic colony and kept at 30°C and 50% RH prior to injection. To prevent cross-contamination, the injection trials were conducted separately for each virus (first using IAPV inoculum and the second with DWV inoculum) three weeks apart. Each trial contained six treatments: untreated pupae, and one microliter injected pupae with either PBS as a mock control or virus solution at different concentrations diluted in PBS. The virus concentrations were low (10 viral genomic copy number of IAPV or DWV diluted inoculum), medium-low (10^2^ viral genomic copy number of IAPV or DWV diluted inoculum), medium-high (10^5^ or 10^4^ viral genomic copy number of IAPV or DWV diluted inoculum, respectively), and high (10^9^ or 10^7^ viral genomic copy number of IAPV or DWV undiluted inoculum, respectively). Injection was conducted manually using a Hamilton ten-microliter syringe. The needle was inserted between the 2^nd^ and 3^rd^ integuments of the pupa’s abdomen. A total of 20 pupae/treatment/virus were placed in a petri dish with filter paper, and incubated under controlled conditions (34 °C, 70% RH). The mortality and developmental stage of the pupae were evaluated daily according to the color of body parts (74); dead pupae were recorded and then discarded. The proportion of pupae either dead or developing in each group over the five days was calculated on the last day. After five days of incubation at least five pupae per treatment were sacrificed for further analysis of chemical cues, viral loads and expression of immune genes. Each pupa was first washed in 0.5mL of hexane for 10 min for the extraction of chemical cues. After hexane removal, the pupae were frozen at - 80°C till further RNA extraction.

### 4.5 Chemical cues extraction

The hexane extracts of five individual pupae for each treatment and ten for each PBS and control were spiked with 10 μL (1ng/μL) of internal standard (butyl butyrate) as in Wagoner et al. (2019) (24) and concentrated to approximately 20 μL under gentle nitrogen flow. For each sample, 1 μL of the extract was analyzed by Trace GC-FID, Thermo Finnigan, USA, using ZB-5, 30m x 0.25mm x 0.25 mm capillary column, and helium as carrier gas at 1mL/min. The detector was set to 270°C and the inlet to 280°C. The oven temperature was programmed as follows: the initial oven temperature was set at 60°C for 2 minutes, followed by temperature increase up to 280°C at 10°C/min. The chromatogram integration was initially conducted on 155 consistent peaks using Chromeleon™ Chromatography Data System. The relative quantity of each selected peak was calculated and subsequently used for statistical analysis.

The chemical profile was further analysed on pools of each experimental group using a GC–MS 7890B/5977A quadrupole, with electron impact ionization (Agilent Technologies, Santa Clara, CA, USA using Agilent HP-5MS without man (30 m × 0.25 mm, 0.25 μm) using helium as the carrier gas at a flow rate of 1mL/min. The injector temperature was the same as the GC. The mass spectrometry was operated at a 70-eV ionization voltage. Peak discrimination was carried out using the MSD Chemstation software (Version F.01.03.2357, Agilent Technologies, Santa Clara, CA, USA). Subsequently, the components were identified according to their fragmentation patterns, published spectra a set of external hydrocarbon standards (from C14 to C33) and by comparison with the NIST/EPA/NIH Mass Spectral Library (Version 2.0f, Agilent Technologies, Santa Clara, CA, USA).

### 4.6 RNA Isolation and cDNA preparation for PCR

Total RNA was extracted from individual pupae using Tri-reagent, as described by Zioni et al. (2011) (42) with some modifications. cDNA was prepared using RevertAid Reverse Transcriptase (Thermo Fisher Scientific Waltham, MA, USA) with oligo-dT and random primers according to the manufacturer’s instructions. RNA quality of individual pupae was assessed via a Nanodrop 2000 spectrophotometer (Thermo Fisher). Five hundred nanograms of RNA template were used to synthesize cDNA. For reverse transcription, RNA and primers were incubated at 65°C for 5 min, followed by the addition of buffer containing 50 mM of Tris-HCl (pH 8.3), 75 mM of KCl, 2 mM of MgCl2, 5 mM of DTT, 4 units of RNase inhibitor Ribolock® (Thermo Fisher Scientific Waltham, MA, USA), and the reverse transcriptase (200 units; Thermo Fisher Scientific Waltham, MA, USA) in a 20 µL volume, and further incubation at 45°C for 45 min. The reaction was terminated by heating at 70°C for 10 min.

### 4.7 Viral loads and immune gene expression

Quantitative PCR was used to assess viral loads (DWV and IAPV) and the relative expression of immune (*PGRP-S2*, *hymenoptaecin*, *Pl2*, *vago*, *abaecin*, *defensin-1*, lysozyme-1) genes using *RPL8* as a reference gene and primers, as described in Table 2S. Viral loads and immune genes were tested in individual pupae. Two technical replicates were performed for each sample. Non-template controls (water) were included for each assay. A 40-cycle reaction was performed on an Applied Biosystems StepOne Plus qPCR machine with the following conditions: pre-incubation 95 ◦C for 2 mins of 95 ◦C for 10 s, 60 ◦C for 20 s, and 72 ◦C for 20 s, with a final melt curve analysis at 65 ◦C for 5 s to 95 ◦C. Viral loads were calculated using a standard curve, as described in Erez et al. (2022). The relative expression levels of genes were calculated using the 2^-ΔCT^ method, as previously published (75).

### 4.8 Statistical analysis

Statistical analysis was performed using JMP 17.0. The percentage of developed pupae was calculated using: the developmental stage of the most developed untreated pupae on day five, as a reference. Subsequently, we calculated the percentage of pupae that reached this stage in each treatment group. Percentage of dead pupae out of the total pupae, accumulated till day five, was calculated for each group. The values of immune gene expression and viral loads (log-transformed) were examined for normality (Shapiro-Wilk test) and homoscedasticity (Levene test). As both tests were significant for all variables, meaning the data were not normally distributed, nor homoscedastic, we performed rank transformations, and the models were rerun on the transformed data. The model considered was a two-way ANOVA on the treatment effect (high, medium-high, medium-low, low, PBS and untreated) and the virus injection effect (DWV/IAPV). Pearson correlation was used to assess the association between viral loads, and the expression of immune genes, in IAPV-infected pupae, DWV-infected pupae, PBS-injected pupae and untreated. For this analysis we grouped all IAPV-injected treatments (low, medium-low, medium-high and high), DWV-injected treatments (low, medium-low, medium-high and high), PBS-injected and untreated pupae of both trials. Statistical analysis on pupae volatiles was conducted on both experiments together, since both trials were conducted on pupae from the same H colony and the profiles of control groups were similar, even though the experiments were conducted three weeks apart. The analysis was done on the relative quantity of all clear 155 GC peaks. These were compared among all treatment groups by one-way ANOVA, followed by post-hoc pairwise comparisons followed by the Bonferroni correction (p<0.0003). This analysis revealed 16 peaks whose relative quantity was significantly different among the treatments. Subsequently, we conducted principal component analysis using these 16 peaks.

## Acknowledgments

We wish to thank: Prof. Abraham Hefetz for his assistance with MS component identifications, Dr. Hillary Voet for her help with statistical analyses, and BARD Binational Foundation grant IS-5469-22 to VS, DS and MS for funding this research.

## Author Contributions

**Conceptualization:** Tal Erez, Victoria Soroker, Nor Chejanovsky, Marla Spivak, Declan C. Schroeder.

**Data Curation**: Tal Erez, Angelina Fathia Osabutey, Elad Bonda, Assaf Otmy, Sofia Levin-Nikulin, Poppy J Hesketh-Best, Clarissa Pellegrini Ferreira, Declan C. Schroeder, Victoria Soroker.

**Formal Analysis:** Tal Erez, Angelina Fathia Osabutey, Elad Bonda, Poppy J Hesketh-Best, Clarissa Pellegrini Ferreira, Declan C. Schroeder, Victoria Soroker.

**Funding acquisition:** Victoria Soroker, Marla Spivak, Declan C. Schroeder.

**Investigation:** Tal Erez, Angelina Fathia Osabutey, Elad Bonda, Assaf Otmy, Poppy J Hesketh-Best, Clarissa Pellegrini Ferreira, Declan C. Schroeder, Victoria Soroker.

**Methodology:** Tal Erez, Sofia Levin-Nikulin, Angelina Fathia Osabutey, Victoria Soroker, Nor Chejanovsky, Marla Spivak, Declan C. Schroeder.

**Project Administration:** Victoria Soroker, Marla Spivak, Declan C. Schroeder.

**Resources:** Victoria Soroker, Declan C. Schroeder.

**Supervision:** Victoria Soroker, Nor Chejanovsky, Marla Spivak, Declan C. Schroeder.

**Validation:** Victoria Soroker, Marla Spivak, Declan C. Schroeder.

**Visualization:** Tal Erez, Angelina Fathia Osabutey, Poppy J Hesketh-Best, Clarissa Pellegrini Ferreira, Declan C. Schroeder, Victoria Soroker.

**Writing – Original Draft Preparation:** Tal Erez.

**Writing – Review & Editing**: All authors.

