## Supplementary for "The effect of two distinct viral-enriched inocula on the immune response and chemical cues in honey bee pupae"

**Table 1S. Summary of viral contigs (and draft/complete genomes) binned from the experimental colony.** Outside parenthesis is the number of contigs, and inside is the number of validated draft/complete genomes.

| Tube ID | Description | Bin_1_DWV contigs (min 8kb genomes) | Bin_2_SBV contigs (min 6kb genomes) | Bin_3_IAPV contigs (min 6kb genomes) | Bin_4_LSV contigs (min 3.5kb genomes) |
|---|---|---|---|---|---|
| **108** | Hive RNA | 1 (0) | 0 (0) | 0 (0) | 1 (1) |

**Table 2S. Oligonucleotide primers sequence tested in this study.**

| Gene/Virus | Sequence | References | |
|---|---|---|---|
| DWV-A | TGGCTAACCGTCGTAAGGCG<br>TAACTGACGCACTAATTTCCGC | Evans 2006; Hou et al. 2014; Zioni et al. 2011; Erez et al. 2022 | |
| DWV-B | TGGCTAATCGACGTAAAGCA<br>ACTAATCTCTGAGCCAACACGT | Evans 2006; Hou et al. 2014; Zioni et al. 2011; Erez et al. 2022 | |
| IAPV | CCAGCCGTGAAACATGTTCTTACC<br>ACATAGTTGCACGCCAATACGAGAAC | Evans 2006; Hou et al. 2014; Zioni et al. 2011; Erez et al. 2022 | |
| *RLP8* | TGGATGTTCAACAGGGTTCATA<br>CTGGTGGTGGACGTATTGATAA | Evans 2006; Hou et al. 2014; Zioni et al. 2011; Erez et al. 2022 | GB17629 |
| *PGRPS-2* | TAATTCATCATTCGGCGACA<br>TGTTTGTCCCATCCTCTTCC | Evans et al. 2006 | GB19301 |
| *Hymenoptaecin* | ACAATGGATTATATCCCGACTCGT<br>CAATGTCCAAGGATGGACGAC | Hinshaw et al. 2021 | GB17538 |

| | | | |
|---|---|---|---|
| *Protein lethal 2 (Pl2)* | ATTTGGATCGTCCACATCGT<br>CGGACAATGGCCGATAGTAG | Brutscher et al. 2017 | GB10397 |
| *Abaecin* | CAGCATTCGCATACGTACCA<br>GACCAGGAAACGTTGGAAAC | Evans et al. 2006 | GB18323 |
| *Vago* | TTTTCGCTGCCGAGGAGAAG<br>GCACATACCGGGAAAATCGC | Maori et al. 2019 | GB10896 |
| *Defensin-1* | GTTGAGGATGAATTCGAGCC<br>TTAACCGAAACGTTTGTCCC | Aronstein et al., 2010 | GB19392; GB41428 |
| *Lysozyme-1* | GGAGGCGAGGATTCTGACTCAATC<br>TGTTGCATATCCCTCCGCTGTG | Aronstein et al., 2010 | XM_001120995 |

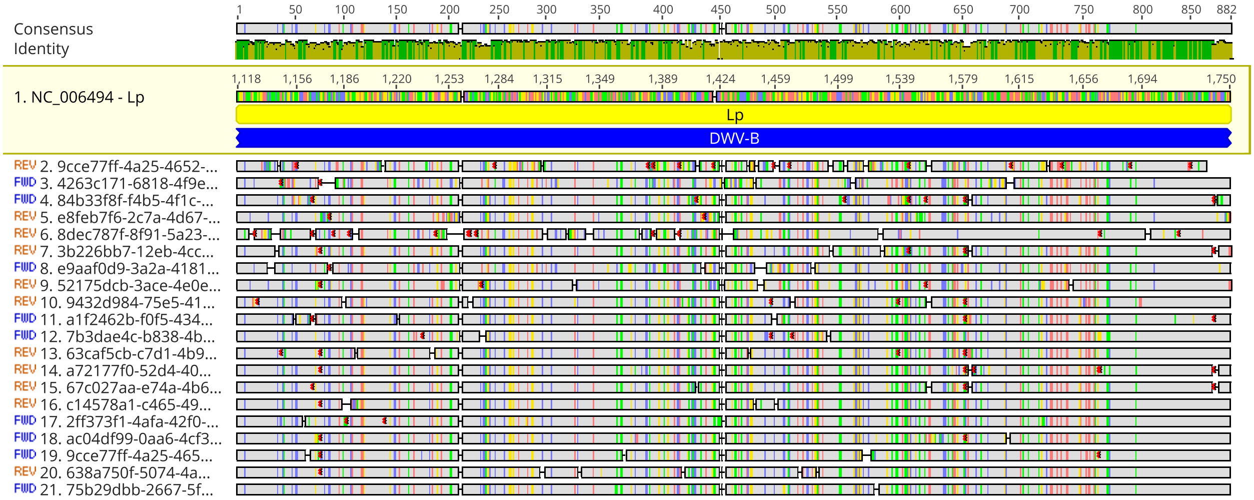


**Fig. 1S.** Read alignment to the Deformed Wing Virus B (DWV-B) genome (NC_006494) in the Lp region from DWV-B/A recombinant samples collected in Minnesota. Each row represents a single read, and mismatches or indels are marked. This recombinant was identified in the DWV inoculum used in the experiment.

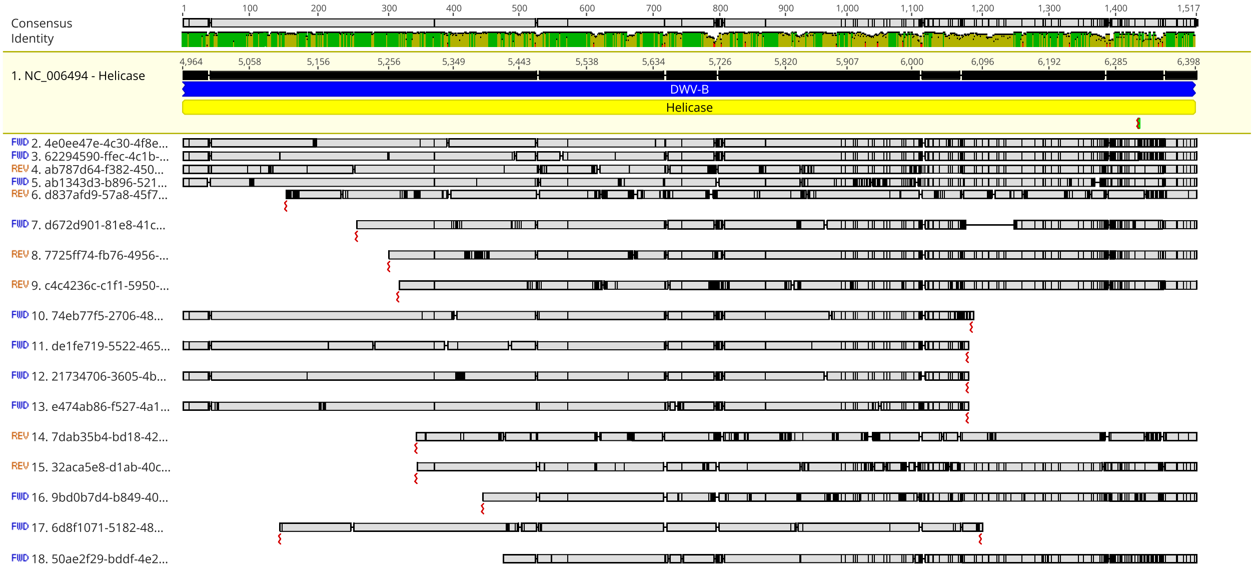


**Fig. 2S.** Read alignment to the Deformed Wing Virus B (DWV-B) genome (NC_006494) in the Helicase region from DWV-B/A recombinant samples collected in Minnesota. Each row represents a single read, and mismatches or indels are marked. This recombinant was identified in the DWV inoculum used in the experiment.

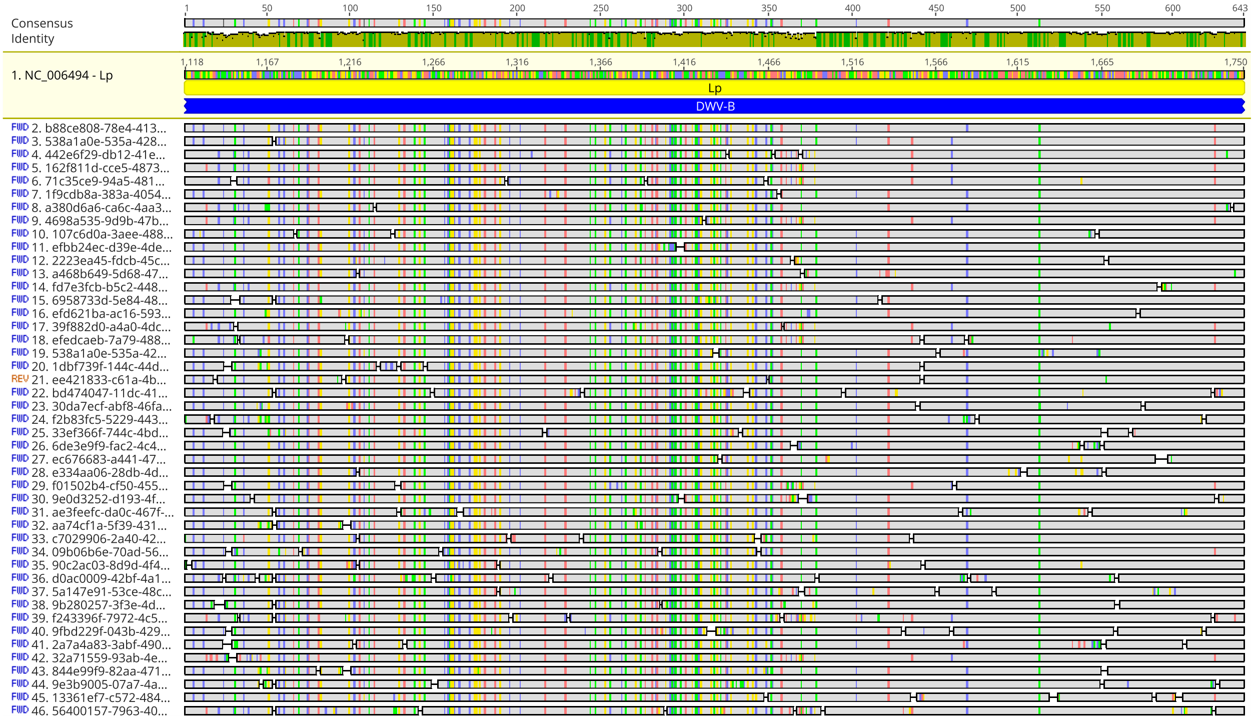


**Fig. 3S.** Read alignment to the Deformed Wing Virus B (DWV-B) genome (NC_006494) in the Lp region from DWV-B/A recombinant samples collected in Israel. Each row represents a single read, and mismatches or indels are marked. This recombinant was identified in the IAPV inoculum used in the experiment.

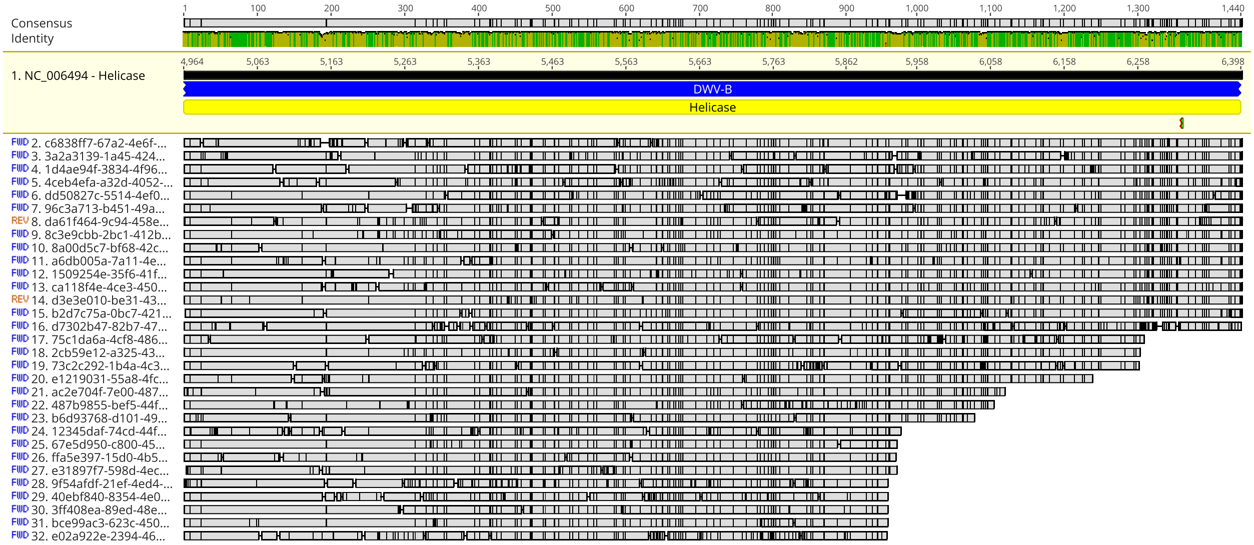


**Fig. 4S.** Read alignment to the Deformed Wing Virus B (DWV-B) genome (NC_006494) in the Helicase region from DWV-B/A recombinant samples collected in Israel. Each row represents a single read, and mismatches or indels are marked. This recombinant was identified in the IAPV inoculum used in the experiment.

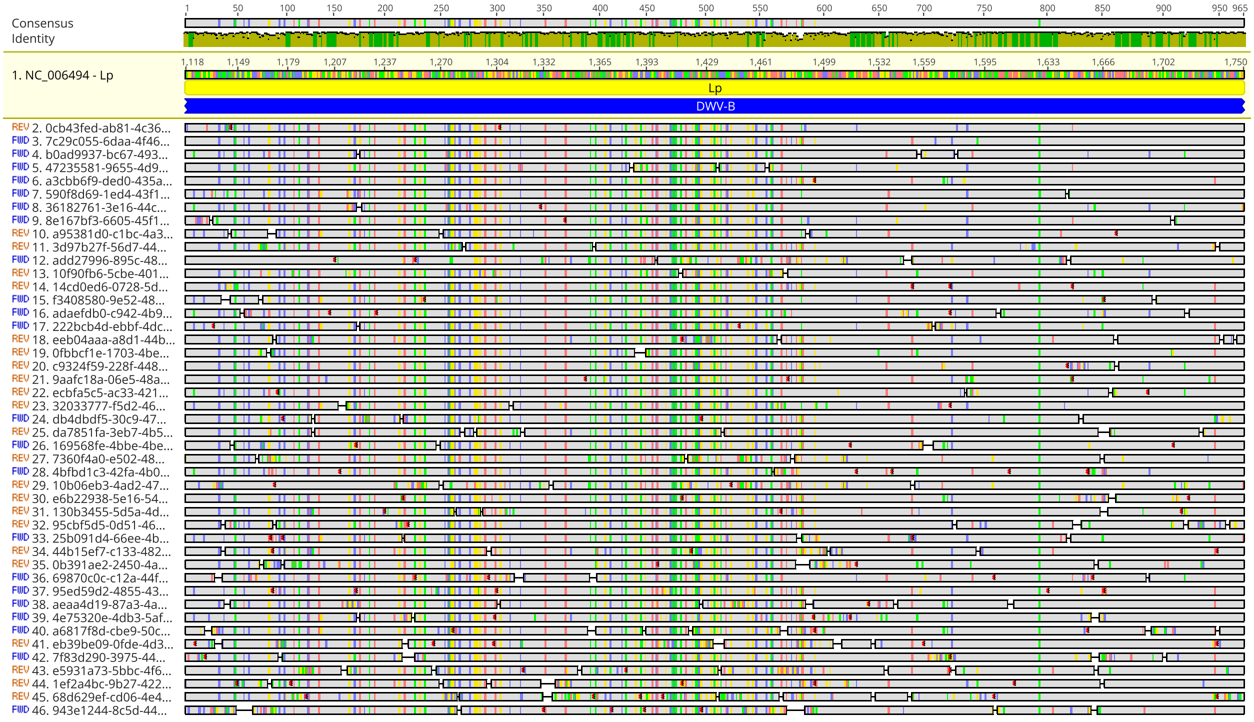


**Fig. 5S.** Read alignment to the Deformed Wing Virus B (DWV-B) genome (NC_006494) in the Lp region from DWV-B/A recombinant samples collected in Israel. Each row represents a single read, and mismatches or indels are marked. This recombinant was identified in the DWV inoculum used in the experiment.

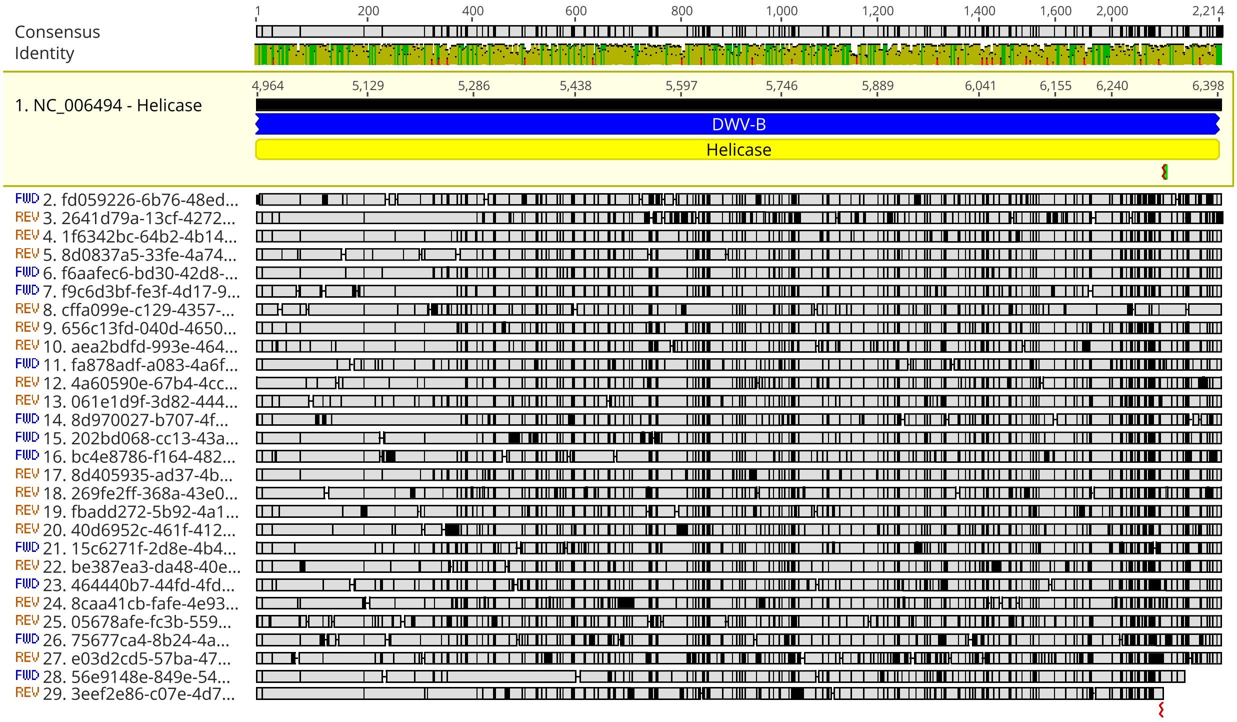


**Fig. 6S.** Read alignment to the Deformed Wing Virus B (DWV-B) genome (NC_006494) in the Helicase region from DWV-B/A recombinant samples collected in Israel. Each row represents a single read, and mismatches or indels are marked. This recombinant was identified in the DWV inoculum used in the experiment.

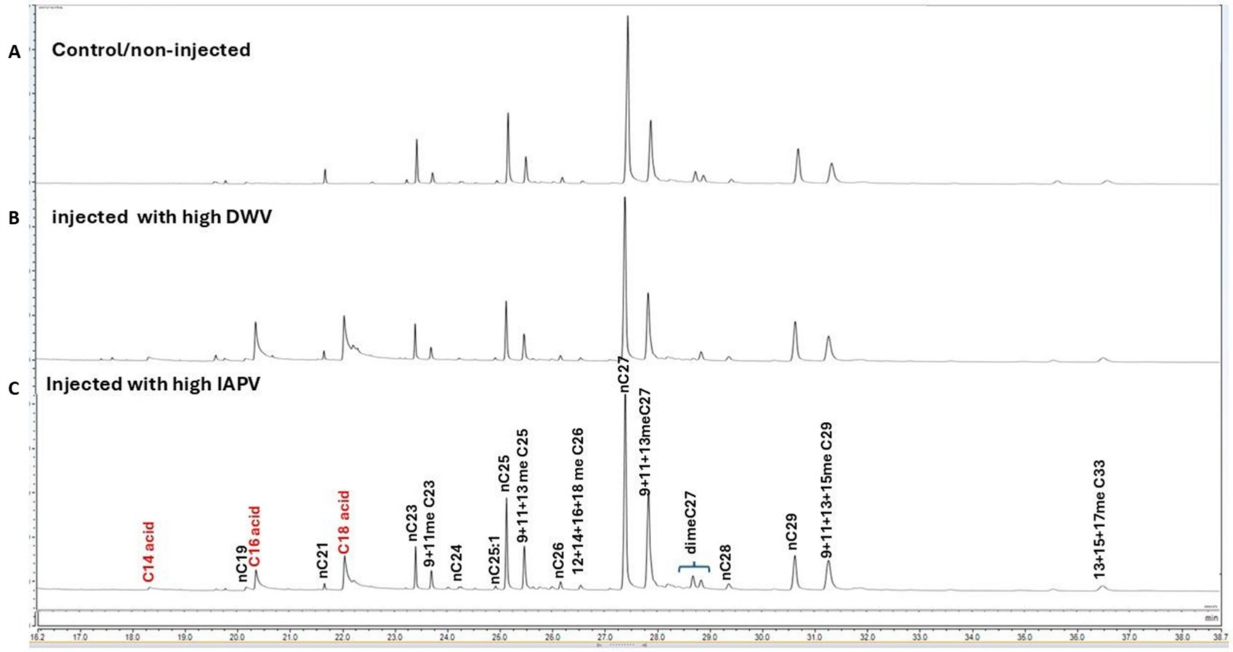

**Fig. 7S.** GC profile of hexane extracts from pools of 10 pupae from non-injected control (A) and two treatment groups: injected with high concentration ($10^6$ copy numbers) of DWV recombinant (B), and (C) injected with high concentration of IAPV ($10^9$ copy numbers).

**Table 3S. Summary of volatiles identified in the pooled extract by GCMS and additional unidentified peaks.** The data are average of relative quantity of five samples ± SE. The colored compounds represent the 20 peaks whose relative quantity is significantly different between treatments (One-way ANOVA, $p<0.05$, followed by Tukey-Kramer HSD).

| Component | Name | Untreated | PBS | IAPV low | DWV low | IAPV high | DWV high |
|---|---|---|---|---|---|---|---|
| Alkanes | Tetradecane | 0.04±0.003 | 0.04±0.003 | 0.05±0.003 | 0.03±0.006 | 0.04±0.005 | 0.04±0.003 |
| | Pentadecane | 0.04±0.005 | 0.04±0.004 | 0.07±0.005 | 0.06±0.009 | 0.06±0.009 | 0.05±0.007 |
| | Hexadecane | 0.04±0.004 | 0.04±0.002 | 0.05±0.006 | 0.05±0.007 | 0.06±0.009 | 0.044±0.004 |
| | Heptadecane | 0.03±0.004<br>C | 0.02±0.002<br>C | 0.09±0.015<br>B | 0.09±0.008<br>B | 0.05±0.004<br>BC | 0.149±0.019<br>A |
| | Octadecane | 0.04±0.003 | 0.04±0.002 | 0.05±0.007 | 0.04±0.006 | 0.05±0.008 | 0.04±0.004 |
| | Nonadecane | 0.40±0.063 | 0.35±0.048 | 0.18±0.010 | 0.25±0.033 | 0.16±0.018 | 0.23±0.036 |
| | Eicosane | 0.08±0.007 | 0.07±0.005 | 0.46±0.117 | 0.15±0.039 | 0.15±0.029 | 0.23±0.065 |
| | Heneicosane | 1.92±0.332<br>A | 1.56±0.220<br>A | 0.28±0.023<br>B | 0.53±0.057<br>B | 0.38±0.043<br>B | 0.53±0.074<br>B |
| | Docosane | 0.23±0.023 | 0.16±0.016 | 0.66±0.147 | 0.26±0.041 | 0.31±0.062 | 0.56±0.152 |
| | Tricosane | 5.81±0.935<br>A | 4.37±0.601<br>AB | 2.05±0.095<br>B | 2.32±0.204<br>B | 2.52±0.039<br>B | 2.22±0.279<br>B |
| | Tetracosane | 0.18±0.094 | 0.08±0.034 | 0.04±0.007 | 0.02±0.002 | 0.05±0.013 | 0.01±0.001 |
| | Pentacosane | 6.45±0.295 | 6.15±0.373 | 5.55±0.273 | 5.82±0.320 | 6.57±0.347 | 5.02±0.217 |
| | Hexacosane | 0.50±0.036 | 0.54±0.032 | 0.46±0.032 | 0.52±0.046 | 0.52±0.071 | 0.45±0.029 |
| | Heptacosane | 22.38±1.305 | 22.07±1.319 | 19.08±1.209 | 23.60±1.303 | 20.53±2.199 | 21.50±0.714 |
| | Octacosane | 2.08±0.740 | 1.48±0.539 | 1.65±0.295 | 0.29±0.028 | 1.88±0.187 | 0.34±0.029 |
| | Nonacosane | 7.85±0.910 | 8.10±0.968 | 5.90±0.655 | 10.02±0.659 | 5.89±1.164 | 9.04±1.010 |
| | Triacontane | 0.14±0.012 | 0.15±0.011 | 0.13±0.011 | 0.19±0.022 | 0.13±0.023 | 0.17±0.025 |
| | Hentriacontane | 0.16±0.038 | 0.90±0.010 | 0.09±0.013 | 0.20±0.101 | 0.11±0.018 | 0.10±0.007 |
| | Tritriacontane | 0.32±0.076 | 0.16±0.027 | 0.17±0.019 | 0.36±0.189 | 0.14±0.019 | 0.14±0.011 |
| Monomethyl alkanes | 9-monomethyl tricosane | 0.03±0.002 | 0.03±0.001 | 0.02±0.002 | 0.03±0.005 | 0.02±0.004 | 0.03±0.005 |
| | 11+13 monomethyl tricosane | 1.20±0.077 | 1.14±0.065 | 1.06±0.090 | 1.05±0.102 | 1.32±0.139 | 0.98±0.097 |
| | 12-monomethyl tetracosane | 0.07±0.007 | 0.06±0.006 | 0.06±0.005 | 0.06±0.007 | 0.06±0.006 | 0.04±0.004 |
| | 5-monomethyl pentacosane | 0.24±0.025<br>B | 0.25±0.041<br>B | 0.29±0.053<br>AB | 0.18±0.023<br>AB | 0.32±0.092<br>B | 0.14±0.013<br>B |

| | | | | | | | |
|---|---|---|---|---|---|---|---|
| | **7-monomethyl pentacosane** | **0.19±0.012** | **0.20±0.009** | **0.18±0.020** | **0.21±0.016** | **0.21±0.022** | **0.17±0.007** |
| | **11+13 monomethyl pentacosane** | **3.24±0.245** | **3.24±0.183** | **3.05±0.221** | **3.04±0.296** | **3.56±0.372** | **2.68±0.085** |
| | **12+14+16+18 monomethyl hexacosane** | **0.09±0.018** | **0.07±0.008** | **0.04±0.006** | **0.06±0.009** | **0.04±0.008** | **0.05±0.007** |
| | **5+7 monomethyl heptacosane** | **0.33±0.041** | **0.38±0.036** | **0.48±0.102** | **0.40±0.053** | **0.34±0.042** | **0.37±0.022** |
| | **9+11+13-monomethyl heptacosane** | **12.52±0.968** | **13.70±0.728** | **12.58±0.754** | **13.43±1.038** | **13.61±1.339** | **11.71±0.682** |
| | **14+16+18 monomethyl octacosane** | **1.61±0.088** | **1.55±0.107** | **1.45±0.100** | **1.72±0.116** | **1.44±0.047** | **1.63±0.103** |
| | **9+11monomethyl nonacosane** | **7.10±0.596** | **8.14±0.373** | **7.23±0.436** | **8.72±0.544** | **7.49±0.841** | **7.78±0.743** |
| | **13+15-monomethyl nonacosane** | **0.41±0.041** | **0.45±0.032** | **0.45±0.057** | **0.51±0.050** | **0.38±0.076** | **0.45±0.055** |
| | **12+14+16-monomethyl triacontane** | **0.19±0.018** | **0.18±0.012** | **0.14±0.008** | **0.22±0.016** | **0.15±0.016** | **0.18±0.019** |
| | **11+13+15-monomethyl hentriacontane** | **1.14±0.285** | **0.72±0.086** | **0.55±0.058** | **1.26±0.400** | **0.57±0.097** | **0.83±0125** |
| | **14-monomethyl dotriacontane** | **0.04±0.004 B** | **0.03±0.003 B** | **0.07±0.008 A** | **0.06±0.010 AB** | **0.05±0.007 AB** | **0.05±0.006 AB** |
| | **18-monomethyl dotriacontane** | **0.13±0.006** | **0.14±0.010** | **0.11±0.007** | **0.14±0.011** | **0.12±0.024** | **0.11±0.012** |
| | **13+15+1-monomethyl tritriacontane** | **0.53±0.043** | **0.62±0.040** | **0.53±0.043** | **0.70±0.056** | **0.55±0.102** | **0.56±0.061** |
| **Dimethyl alkanes** | **7, 13-dimethyl pentacosane** | **0.11±0.011** | **0.10±0.012** | **0.11±0.019** | **0.09±0.012** | **0.12±0.035** | **0.09±0.009** |
| | **9, 13-dimethyl pentacosane** | **0.24±0.021** | **0.22±0.026** | **0.24±0.025** | **0.19±0.025** | **0.25±0.055** | **0.15±0.018** |
| | **5, 15-dimethyl pentacosane** | **0.09±0.022** | **0.1±0.015** | **0.09±0.010** | **0.07±0.006** | **0.09±0.026** | **0.07±0.013** |
| | **5,15-dimethyl heptacosane** | **0.76±0.083** | **0.84±0.062** | **0.85±0.086** | **0.78±0.097** | **0.81±0.137** | **0.71±0.076** |

| | | | | | | | |
|---|---|---|---|---|---|---|---|
| | 5, 17-+ 7,13 +11,15 dimethyl heptacosane | 0.35±0.042 | 0.37±0.033 | 0.45±0.080 | 0.36±0.047 | 0.36±0.073 | 0.32±0.028 |
| | 7, 17-dimethyl heptacosane | 0.38±0.049 | 0.42±0.037 | 0.50±0.058 | 0.44±0.057 | 0.40±0.076 | 0.39±0.049 |
| | 9, X+14,16 -dimethyl octacosane | 0.92±0.077 | 1.02±0.061 | 0.94±0.060 | 1.00±0.076 | 1.00±0.144 | 0.83±0.090 |
| | 16,18-dimethyl octacosane | 0.15±0.017 | 0.13±0.017 | 0.22±0.101 | 0.14±0.005 | 0.15±0.045 | 0.14±0.026 |
| | 13, 15-dimethyl nonacosane | 0.15±0.017 | 0.13±0.017 | 0.22±0.101 | 0.14±0.005 | 0.15±0.045 | 0.14±0.026 |
| | 7, 15+ 9,15+ 11, 15-dimethyl nonacosane | 0.78±0.079 | 0.81±0.044 | 0.81±0.078 | 1.00±0.104 | 0.76±0.147 | 0.87±0.128 |
| | 5,15 dimethyl hentriacontane | 2.05±0.179 | 2.36±0.120 | 2.07±0.145 | 2.63±0.172 | 2.15±0.365 | 2.37±0.258 |
| | 5,17 dimethyl hentriacontane | 0.25±0.019 | 0.29±0.018 | 0.27±0.020 | 0.31±0.024 | 0.26±0.047 | 0.25±0.033 |
| | 13,17+13,19 dimethyl hentriacontane | 0.07±0.005 | 0.06±0.004 | 0.06±0.003 | 0.07±0.011 | 0.07±0.010 | 0.06±0.006 |
| | 11,19 dimethyl hentriacontane | 0.04±0.006 | 0.04±0.006 | 0.04±0.010 | 0.05±0.010 | 0.03±0.006 | 0.03±0.008 |
| Alkenes | Heneicosene | 0.10±0.019 | 0.07±0.009 | 0.08±0.010 | 0.12±0.051 | 0.09±0.007 | 0.13±0.039 |
| | Tricosene | 0.88±0.210 A | 0.54±0.125 AB | 0.18±0.065 B | 0.11±0.028 B | 0.10±0.007 B | 0.17±0.047 B |
| | Pentacosene | 0.53±0.083 A | 0.43±0.041 AB | 0.22±0.031 C | 0.27±0.021 BC | 0.31±0.020 ABC | 0.25±0.019 BC |
| | Heptacosene | 0.12±0.016 | 0.11±0.005 | 0.10±0.008 | 0.15±0.026 | 0.11±0.009 | 0.12±0.012 |
| | Octacosene | 0.26±0.041 | 0.29±0.031 | 0.37±0.056 | 0.31±0.038 | 0.26±0.061 | 0.26±0.040 |
| | Nonacosene | 0.28±0.016 | 0.30±0.015 | 0.23±0.020 | 0.32±0.036 | 0.29±0.033 | 0.32±0.026 |
| | Hentriacontene | 0.29±0.023 | 0.28±0.021 | 0.23±0.020 | 0.37±0.057 | 0.25±0.038 | 0.27±0.029 |
| | Tritriacontene | 0.06±0.008 | 0.06±0.003 | 0.05±0.005 | 0.06±0.003 | 0.05±0.017 | 0.05±0.003 |
| Fatty acids | Nonanoic acid | 0.06±0.008 | 0.05±0.007 | 0.05±0.006 | 0.05±0.004 | 0.05±0.011 | 0.04±0.009 |
| | Decanoic acid | 0.11±0.015 | 0.112±0.007 | 0.111±0.011 | 0.108±0.017 | 0.102±0.016 | 0.083±0.012 |
| | Dodecanoic | 0.03±0.003 | 0.03±0.002 | 0.10±0.062 | 0.03±0.004 | 0.04±0.004 | 0.02±0.003 |

| | | | | | | | |
|---|---|---|---|---|---|---|---|
| | Tetradecanoic acid | 0.02±0.002 | 0.01±0.002 | 0.03±0.005 | 0.02±0.003 | 0.02±0.005 | 0.02±0.004 |
| | Hexadecanoic acid | 0.20±0.031 B | 0.19±0.020 B | 3.29±0.758 A | 0.81±0.266 B | 1.04±0.282 B | 1.49±0.426 AB |
| | Octadecanoic acid | 0.08±0.006 C | 0.09±0.010 C | 4.58±0.889 A | 1.49±0.514 BC | 2.01±0.660 ABC | 2.56±0.347 AB |
| Esters | Ethyl oleate | 0.06±0.016 C | 0.05±0.009 C | 2.53±0.0494 A | 0.73±0.205 BC | 1.39±0.435 ABC | 1.79±0.587 AB |
| | Octadecenoic acid, ethyl ester | 0.04±0.004 | 0.06±0.026 | 0.63±0.203 | 0.23±0.077 | 0.27±0.085 | 0.42±0.094 |
| | Hexanedioic acid, dioctyl ester | 0.04±0.009 | 0.04±0.008 | 0.04±0.007 | 0.02±0.008 | 0.03±0.011 | 0.03±0.005 |
| | Isopropyl palmitate | 0.06±0.010 | 0.06±0.008 | 0.17±0.029 | 0.06±0.009 | 0.11±0.019 | 0.06±0.014 |
| unknown significant peaks | unknown | 0.07±0.017 B | 0.07±0.017 B | 0.09±0.020 B | 0.10±0.013 B | 0.10±0.025 B | 0.12±0.013 A |
| | unknown | 0.01±0.002 B | 0.01±0.002 B | 0.01±0.001 A | 0.07±0.006 A | 0.07±0.016 A | 0.06±0.009 A |
| | unknown | 0.02±0.002 A | 0.02±0.002 AB | 0.01±0.001 B | 0.01±0.002 A | 0.02±0.002 B | 0.01±0.001 AB |
| | unknown | 0.22±0.033 A | 0.22±0.033 AB | 0.17±0.026 C | 0.04±0.021 C | 0.01±0.002 BC | 0.05±0.034 C |
| | unknown | 0.02±0.002 B | 0.02±0.002 B | 0.02±0.002 AB | 0.04±0.010 AB | 0.03±0.003 AB | 0.05±0.011 A |
| | unknown | 0.10±0.018 A | 0.10±0.018 A | 0.08±0.014 B | 0.02±0.004 B | 0.01±0.004 B | 0.01±0.002 AB |
| | unknown | 0.15±0.012 B | 0.15±0.012 B | 0.09±0.015 A | 1.94±0.641 AB | 0.73±0.294 AB | 0.55±0.127 AB |
| | unknown | 0.04±0.002 B | 0.04±0.002 B | 0.03±0.003 B | 0.03±0.002 BC | 0.04±0.005 A | 0.04±0.005 C |
